# Microbial Named Entity Recognition and Normalisation for AI-assisted Literature Review and Meta-Analysis

**DOI:** 10.1101/2025.08.29.671515

**Authors:** Dhylan Patel, Antoine D. Lain, Avish Vijayaraghavan, Nazanin Faghih Mirzaei, Monica N. Mweetwa, Meiqi Wang, Tim Beck, Joram M. Posma

## Abstract

**Motivation:** Manual curation of biomedical literature is slow and error-prone and while large language models (LLMs) trained on general texts have shown to be useful for text summarisation, these methods lack the domain-specific expertise required to perform this task accurately. Here we describe the creation of the first microbiome-specific text corpus, use this to train deep learning algorithms for named-entity recognition (NER) and normalisation (NEN), and demonstrate their use to meta-analyse microbiome literature.

**Methods:** We developed an automated pipeline to annotate all mentions of bacteria, archaea, and fungi in 1,410 full-text microbiome articles. We manually annotated (gold-standard) a separate test set of 288 documents. We trained different transformer-based language models for microbiome recognition and normalisation to taxonomic identifiers and evaluate their performance using the precision, recall, F1-score, and accuracy on the test set. The best models were used to automatically annotate all available Open Access, full-text microbiome articles (n=6,927) and identify taxa that are significantly overrepresented across 14 domains.

**Results:** The training and validation set contained a total of 90,150 annotations (both long form and abbreviations). Using the gold-standard test set, with an inter-annotator agreement rate of 99.52% for NER and 88.31% for NEN, the trained models were evaluated and our fine-tuned BioBERT model achieved an F1-score of 96% for NER surpassing a rule- and dictionary-based annotation pipeline (94%). For NEN the accuracy obtained by the deep learning models greatly surpassed that of the pipeline (91% vs 69%). Evaluated across the entire available literature, our models annotate an entire full-text document in only 7 seconds.

**Conclusion:** Our algorithms have near perfect precision and greatly speed up the process of annotating microbes in full-text articles. We demonstrated the capabilities of these methods by analysing the entire available literature and describe the taxa associated with each of the domains in our meta-analysis, and exemplify how these methods can be integrated into literature review workflows improving both the speed and accuracy of results.

**Availability:** All codes and data for automatic annotation, model training, and generation of taxonomic trees visualising the data will be made available following peer review with instructions on how to deploy the model on new texts from https://github.com/omicsNLP/microbELP.

## Introduction

Manual extraction of information from large numbers of publications is time-consuming, labour-intensive and may lead to human errors. With the biomedical literature expanding at an increased rate (1), there is a clear need to develop new methods for specialised tasks; tasks that large language models (LLMs), such as GPT (2), usually lack extensive training data for, given these LLMs are often pre-trained on general texts and therefore lack specific biomedical knowledge to excel in these areas. It has been demonstrated by different surveys that ‘small’ language models (LMs) such as BERT-based models (3–7) outperform LLMs for named entity recognition (NER) (8–10), entity linking (9), text classification (8, 9) and other tasks not relying on text generation. While LLMs generally perform better for question-answering and relation extraction tasks (8, 9).

To use any type of LM (small or large) for literature review, several elements determine their performance, and therefore usefulness. Domain specificity is important as there may be specific vocabularies and terminologies used, and nuances made in (biomedical) scientific literature that are less common in general text. Training or fine-tuning existing models for specific tasks require structured text corpora with enough annotated training examples for the task to allow them to pick up relevant language patterns. Some corpora exist in which organisms are annotated, such as CRAFT (11) (97 full-texts, 7,449 organisms), Linnaeus (12) (100 full-texts, 4,259 organisms), species-800 (13) (800 abstracts, 3,708 organisms), BioNLP-ID (14) (30 full-texts, 3,471 organisms), miRNA corpus (15) (201 abstracts, 546 organisms), and CellFinder (16) (10 full-texts, 438 organisms). Some of these corpora have been used to train LMs that have shown state-of-the-art performance for biomedical NER, such as BioBERT (3), BERN2 (17), and SciBERT (4).

The microbiome is the collection of microorganisms that play crucial roles in biochemical reactions by modulating processes essential for maintaining host health. While these are organisms, the problem of using existing organism recognition models relates both to specificity and size, since not all species included are microbes and only a subset of microbes are included in these corpora. Therefore these corpora, and by extension the models trained on them, are not suitable for microbiome NER as these will inflate the number of wrongly extracted mentions when aiming to target microbiome entities using the organism tag, and since many species are unseen by the training data they also yield plenty of missed ones.

Here, we aim to fill this gap by describing the creation of a microbiome-specific training corpus, a dictionary-based annotation and normalisation workflow (with microbiome-specific rules), construction of an annotated microbiome (bacteria, archaea, fungi) corpus of full-text publications in machine-readable format, and finally contributing four transformer-based LMs trained on these data and evaluated on a manually annotated test set, including two models for NER and two for Named Entity Normalisation (NEN).

## Materials and Methods

### Data - training corpus

PubMed Central was searched on 14/Apr/2020 to generate an initial corpus spanning 5 different microbiome domains (airway (from oral and nasal to lung), faecal (stool), skin (skin tissue), urinary (excluding vaginal) and vaginal (vagina, cervix, placenta)) in humans for model training (see below). The following search query was used for obtaining all ‘faecal microbiome’ articles: *(metagenomics[Abstract] OR metagenomic[Abstract] OR metagenome[Abstract] OR microbiome[Abstract] OR microbiomic[Abstract] OR 16S[Abstract] OR meta-genome[Abstract] OR microbial[Abstract] OR microbes[Abstract] OR metagenomics[Title] OR metagenomic[Title] OR metagenome[Title] OR microbiome[Title] OR microbiomic[Title] OR 16S[Title] OR meta-genome[Title] OR microbial[Title] OR microbes[Title]) AND (faecal[Abstract] OR faeces[Abstract] OR stool[Abstract] OR feces[Abstract] OR fecal[Abstract] OR faecal[Title] OR faeces[Title] OR stool[Title] OR feces[Title] OR fecal[Title]) AND (16S[Abstract] OR rRNA[Abstract] OR sequencing[Abstract] OR shot-gun[Abstract] OR Illumina[Abstract] OR MiSeq[Abstract] OR PCR[Abstract] OR 16S[Title] OR rRNA[Title] OR sequencing[Title] OR shotgun[Title] OR Illumina[Title] OR MiSeq[Title] OR PCR[Title]) AND OR abundant[Abstract] OR diversity[Abstract] OR abundance[Title] OR abundant[Title] OR diversity[Title]) AND (human[Abstract] OR patients[Abstract] OR subjects[Abstract] OR participants[Abstract] OR human[Title] OR patients[Title] OR subjects[Title] OR participants[Title]) NOT (mouse[Abstract] OR mice[Abstract] OR rat[Abstract] OR rats[Abstract] OR dog[Abstract] OR dogs[Abstract] OR animal[Abstract] OR cell culture[Abstract] OR dose[Abstract] OR review[Title] OR proteomics[Abstract] OR diet[Abstract] OR proteomic[Abstract] OR proteome[Abstract] OR transcriptomics[Abstract] OR transcriptomic[Abstract] OR transcriptome[Abstract] OR mouse[Title] OR mice[Title] OR rat[Title] OR rats[Title] OR dog[Title] OR dogs[Title] OR animal[Title] OR cell culture[Title] OR dose[Title] OR proteomics[Title] OR diet[Title] OR proteomic[Title] OR proteome[Title] OR transcriptomics[Title] OR transcriptomic[Title] OR transcriptome[Title]) AND (“01/01/2000”[Publication Date]* : *“3000”[Publication Date])*. Where the grey colour in the query was replaced for each of the domains with synonyms for each of the domains.

The publications were converted from HTML and XML documents, standardised and converted to the machine-readable BioC-JSON format using Auto-CORPus (18). First, Auto-CORPus converts the main text of each publication from HTML or XML to BioC JSON format. In this file, the pipeline splits the publications into different sections using the Information Artifact Ontology (IAO) (19) and in each section, each paragraph forms a single ‘text’-feature. Second, Auto-CORPus transforms tables inside publications to a table-JSON format, and it extracts abbreviations from both the main text and from separate abbreviations sections (if available) within the main text and generates another JSON file for abbreviations with linked full definitions. Here, we use the full-text BioC-JSON files for pre-processing, see more details in 20, and to build a silver-standard annotated corpus for DL models.

### Data - NCBI taxonomy

The NCBI taxonomy data was downloaded on 01/Nov/2021 as separate memory dump (dmp) files. All names (scientific, common, blast, other) were extracted including the child and parent identifiers of each entry. The data was filtered for any node that is an ancestor of either txid2 (bacteria), txid2157 (archaea) or txid4751 (fungi). Several data cleaning steps were performed: main taxonomic names were extracted from the name fields with species as lowest level. Data with lower taxonomic ranks than species were pruned to their species names, with sub-species given the name of the [genus subspecies] pair as ‘species’ name, with species names extracted from the full names associated with the forma, species group and species subgroup ranks. Data without a rank (‘no rank’) was assigned a taxonomic rank based on the (grand)parent and (grand)child nodes with a likely rank inferred from this. This means an entry with ‘no rank’ whose parent is a genus and with (grand)children from taxonomy ranks such as ‘isolate’, ‘strain’, etc. were assigned to ‘species’.

Duplicate names within the same identifier were removed, in addition to the following rules to remove entries: any ‘species’ without a space in the name, ‘species’ where the cleaned species name includes a ‘.’ but not ‘sp.’, any uncommon ending (e.g. ‘-ic’ or ‘-ing’), any entry where the name includes any of the following terms (group, subdivision, symbiont, cluster, biofilm, candidate, candidatus, unclassified, environmental, witches, marine, enrichment), any entry with a country name or country adjective, any entry with less than 4 lowercase letters, any entry where the cleaned name includes a colour, and any entry where the cleaned name includes the source (e.g. soil, sea(water), landfill) or a non-microbial species (e.g. penguin). The final dictionary was saved as a JSON file containing fields original name, cleaned name, taxonomic rank, taxonomy ID, parent ID and kingdom ID and used subsequently for the annotation pipeline. The summary statistics can be found in Table 1.

**Table 1.**
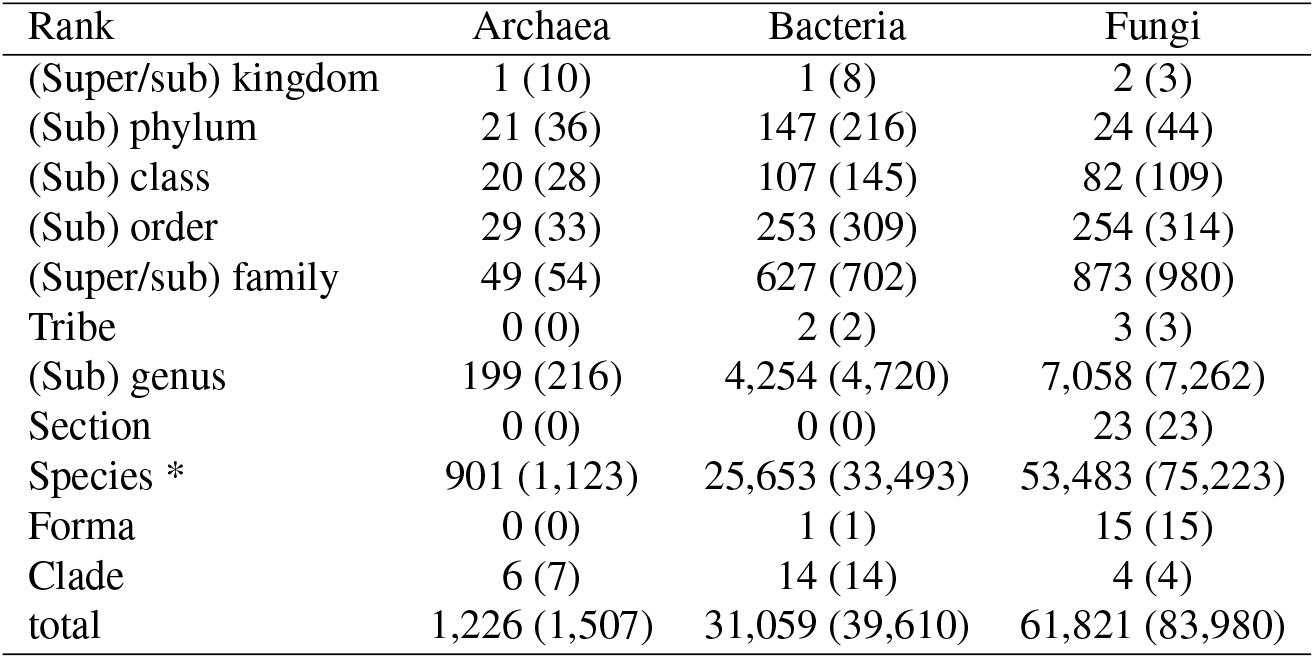
Number of unique identifiers (with number of unique names in brackets) in each of the three microbial kingdoms following data curation. (* includes subspecies and lower insofar these could be mapped to a unique name)

| Rank | Archaea | Bacteria | Fungi |
| --- | --- | --- | --- |
| (Super/sub) kingdom | 1 (10) | 1 (8) | 2 (3) |
| (Sub) phylum | 21 (36) | 147 (216) | 24 (44) |
| (Sub) class | 20 (28) | 107 (145) | 82 (109) |
| (Sub) order | 29 (33) | 253 (309) | 254 (314) |
| (Super/sub) family | 49 (54) | 627 (702) | 873 (980) |
| Tribe | 0 (0) | 2 (2) | 3 (3) |
| (Sub) genus | 199 (216) | 4,254 (4,720) | 7,058 (7,262) |
| Section | 0 (0) | 0 (0) | 23 (23) |
| Species * | 901 (1,123) | 25,653 (33,493) | 53,483 (75,223) |
| Forma | 0 (0) | 1 (1) | 15 (15) |
| Clade | 6 (7) | 14 (14) | 4 (4) |
| total | 1,226 (1,507) | 31,059 (39,610) | 61,821 (83,980) |

### Training and test splits

A total of 1,410 documents were used for training our LMs, with another 288 used for testing both the pipeline and the LMs. The document numbers were derived by splitting roughly 80:20 (train:test) across the 5 domains of articles that were targeted: airway (oral to lung), faeces (gut, stool), skin, urinary and vaginal microbiomes.

A subset of documents in the training portion of the corpus were used to develop the rules implemented in the annotation pipeline (below). The annotated training set (see details on annotations in Table 2) was used as input to the deep learning NER models (below) and split into training and validation portions within the algorithms to fine-tune the models.

**Table 2.**
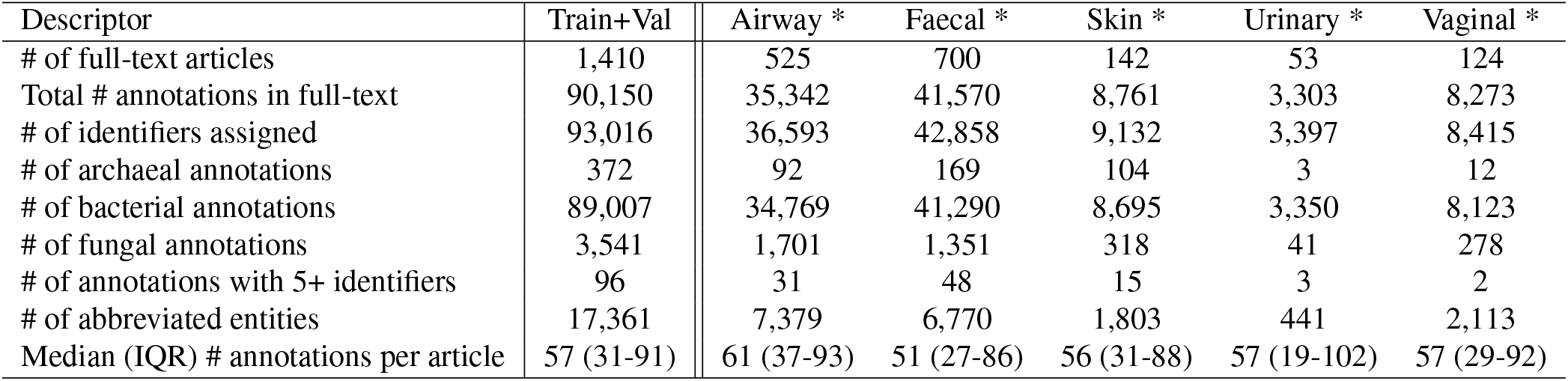
Characteristics of the silver-annotated training+validation corpus. The number of identifiers exceeds the number of annotations as some annotations cannot be resolved to a single identifier due to ambiguity. (* articles can be assigned to multiple domains)

| Descriptor | Train+Val | Airway * | Faecal * | Skin * | Urinary * | Vaginal * |
| --- | --- | --- | --- | --- | --- | --- |
| # of full-text articles | 1,410 | 525 | 700 | 142 | 53 | 124 |
| Total # annotations in full-text | 90,150 | 35,342 | 41,570 | 8,761 | 3,303 | 8,273 |
| # of identifiers assigned | 93,016 | 36,593 | 42,858 | 9,132 | 3,397 | 8,415 |
| # of archaeal annotations | 372 | 92 | 169 | 104 | 3 | 12 |
| # of bacterial annotations | 89,007 | 34,769 | 41,290 | 8,695 | 3,350 | 8,123 |
| # of fungal annotations | 3,541 | 1,701 | 1,351 | 318 | 41 | 278 |
| # of annotations with 5+ identifiers | 96 | 31 | 48 | 15 | 3 | 2 |
| # of abbreviated entities | 17,361 | 7,379 | 6,770 | 1,803 | 441 | 2,113 |
| Median (IQR) # annotations per article | 57 (31-91) | 61 (37-93) | 51 (27-86) | 56 (31-88) | 57 (19-102) | 57 (29-92) |

Each full-text article from the test set (n = 288) was converted to BioC XML and uploaded to a local installation of TeamTat (21) and annotated by two independent annotators resulting in 23,561 microbiome entities and their corresponding identifiers. Our annotators had an inter-annotator agreement of 0.8897 F1-score. Disagreements between pairs of annotators were resolved by a third annotator to arbitrate. All disagreements were resolved through this process, resulting in a minimum of two human annotators having verified each annotation.

### Rule-based annotation pipeline

The (full)text is scanned word for word for any exact matches to words (i.e. the entry *Escherichia coli* is split into two words) in the dictionary. If no match is triggered, the word is converted to Latin singular/plural forms (i.e. *Actinobacterial* to *Actinobacteria*) and checked again. If a match is triggered the words before and after are evaluated. If the word before is already annotated (e.g. a genus) and no punctuation can be found between both words they are combined as a single entity (case for species) to be matched against the names in the dictionary. If the word after triggers a match (without punctuation in between) they are combined and matched to entries in the dictionary. If the word before is a capital letter only, a period was removed after it, and the trigger word is lowercase it is assumed to be an abbreviated entity (e.g. *E. coli*). For abbreviated entities the prior annotations in the document are searched for any match for the full name, if there is a match the abbreviation inherits the ID from the full term, if there is no match then all matches in the dictionary are added (e.g. *M. epidermidis* mapping to *Micrococcus epidermidis* (txid1282) and *Macrococcus epidermidis* (txid1902580)). Once the document is fully annotated it iterates over all annotations with more than one assigned identifier of different taxonomic ranks. For those entries (e.g. *Actinobacteria* phylum and class) the other annotations in the sentence are checked, if only one of the ranks is represented in the other annotations this is the only ID selected for the annotation with more than one. If both or neither are matched, all IDs remain assigned to the entity.

### Deep Learning Models for NER

We used the 1,410 files obtained from our search strategy, already converted from HTML and XML formats to BioC JSON, and ran our pipeline to generate annotations. The annotated BioC JSON files were then converted to the BIO2 (22) format using bins of a maximum size of 512 tokens, stopping at the end of the most recently completed sentence. During this conversion, we excluded the reference sections to reduce potential noise during training due to the formatting of references being different from normal sentences in full text. After conversion to BIO2, we split the data into a 70:30 ratio for training and validation, ensuring the same distribution of the five broad domains used in our search strategy.

We selected two pre-trained models, BioBERT (*biobertbase-cased-v1*.*2*) and SciBERT (*scibert_scivocab_cased*), to fine-tune on our data. For each model, we repeated the process using nine different random seeds to evaluate performance across varying weight initialisations and reduce the influence of luck. An odd number of models was chosen to allow selecting the median model for NEN (see Evaluation Metrics below). We did not perform any hyperparameter search but fine-tuned each model with a fixed set of parameters based on prior work (3, 4): a batch size of 24 bins, a learning rate of 3×10^−5^, five epochs, and a weight decay of 1×10^−5^.

All models were trained on a Linux workstation with an Intel 24-core i9-13900K CPU with 192GB RAM (4×48GB) and an Nvidia GeForce RTX 4090 GPU (24GB memory, 16,384 CUDA cores). The epoch with the highest F1 score (see below) for the validation dataset was stored and used to evaluate on the manually annotated test set. On average the training took 13 minutes per model.

### Deep Learning Models for NEN

To train our deep learning model for NEN, we used the same training and validation sets as those used for NER. The most common approach to training entity representation models involves constructing a pair-wise training dataset. Accordingly, we extracted both the entity and the identifier found by the pipeline when creating the training and validation datasets. We chose to retrain the BioSyn architecture (23) due to its use of embedding space information, its capacity to handle out-of-vocabulary terms, and its ability to perform synonym marginalisation. Similar to the NER task, reference sections were excluded from the datasets.

To retrain a BioSyn model, a dictionary is required on top of the datasets. For this purpose, we used the file generated in the ‘Data - NCBI Taxonomy’ section as the dictionary for training, validation, and testing. Further details regarding the input formatting can be found in the original GitHub repository (https://github.com/dmis-lab/BioSyn).

We used five pre-trained models to develop our BioSyn models. Three models were fine-tuned using their original weights: BioBERT (3), SciBERT (4), and SapBERT (24). The remaining two models were derived from the previous section: the fifth-best (median) fine-tuned BioBERT model and the median fine-tuned SciBERT model, as determined by their performance on the validation set. We did not conduct hyperparameter fine-tuning but instead employed the following fixed parameter set: top-k of 20, 10 epochs, batch size of 16, learning rate of 1×10^−5^, hybrid scoring, and a dense ratio of 0.5.

All models were trained on the same Linux workstation as described previously, with an average training time of 40 minutes per model.

### Comparison of our models for species recognition using BERN2

We compare our microbiome NER results with the BERN2 multi-task model (17), which is based on BioBERT, for the species entity tag. For a quicker turnaround time on our large corpus, we followed the authors’ guidelines in the BERN2 repository (https://github.com/dmis-lab/BERN2) to install and run a local version of the model on the same Linux workstation as above.

Once the BERN2 model is running on a local port within its conda environment, we make requests to it using the URL associated with the port. Running our corpus through the model returns a set of found entities per paper (and optionally per paragraph) which fall into one of nine biomedical classes: gene/protein, disease, drug/chemical, species, mutation, cell line, cell type, DNA, and RNA. We filter entities for the species class to enable microbiome NER and NEN comparison to our models.

### Evaluation Metrics

To evaluate the performance of the annotation pipeline and DL models for NER, we calculate 3 properties of a confusion matrix: true positive (TP) entities, false positives (FPs) and false negatives (FNs). FPs are entities predicted to be a microbial entity by a model which does not match an entity in the test set. FNs are entities that were not annotated by the model but are contained in the manually annotated test set. Using these measures, we calculate the precision (probability of correctly predicted positive values in the total predicted positive outcomes, Equation 1), recall (also known as sensitivity, probability of correctly predicted positives relative to all true positive cases, Equation 2) and F1-score (weighted average of precision and recall, Equation 3).

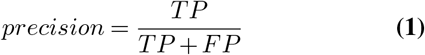

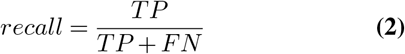

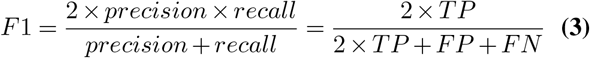

We calculate these metrics according to three methods of evaluating an entity: strict, relaxed and proportional overlap (token based). Strict indicates the exact span of an entity must be found to be counted as TP. A TP for relaxed is counted when the span includes at least one character from the annotation. We define the proportional overlap measure to be an intermediate of the two prior ones, for each annotation the number of tokens correctly identified (proportional to the length of the found entity) is counted towards TP, the proportion of missed tokens to FN and any additional tokens proportionally to FP. The proportional overlap does not penalise partial positive matches, i.e. where only the genus is correctly identified (when part of a species name).

To evaluate the performance of the annotation pipeline and DL models for NEN, we calculate two common accuracy metrics: Acc@1 and Acc@5. These metrics assess the model’s ability to rank the correct identifier among the top candidates predicted for each entity.

**Acc@1** measures the proportion of cases where the top-ranked candidate (rank 1) matches the correct identifier. It is formally defined as:

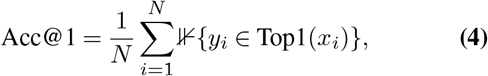

where *N* is the total number of entities, *y*_*i*_ is the correct identifier for the *i*-th entity, *x*_*i*_ is the input entity representation, and ⊮{·} is an indicator function that returns 1 if the condition inside is true, and 0 otherwise. Top1(*x*_*i*_) represents the highest-ranked candidate for *x*_*i*_.

**Acc@5** extends this metric to consider the top five ranked candidates. It is defined as:

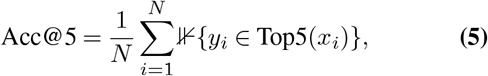

where Top5(*x*_*i*_) represents the set of the five highest-ranked candidates for *x*_*i*_.

These metrics allow us to evaluate how effectively the models rank the correct identifier among the top predictions. Higher Acc@1 indicates the top-ranked prediction is the correct one, while higher Acc@5 indicates the correct identifier is within the top five predictions.

### AI-assisted literature summarisation

The best performing model was applied to all machine-accessible Open Access documents from 14 microbiome domains that were published with Open Access from 01-Jan-2000 to 31-Dec-2023:

oral (mouth and saliva), nasal (including nasopharynx), pharynx (throat), lung (lung, respiratory tract), upper digestive tract (oesophagus to duodenum), lower digestive tract (jejunum to anus), faecal (faecal, stool), hepatobiliary (liver, pancreas, bile duct, gall bladder), skin (skin, toe, foot, elbow fold, forehead), female reproductive (vagina, cervix, endometrium, but not urinary), placental (placenta), breast milk (human milk, colostrum, lactation), male reproductive (testicle, semen, but not urinary), and urinary (urine but not vaginal or testicular) microbiomes. A total of 8,120 articles were returned from the search, with a total of 6,927 available from PubMed Central’s Open Access corpus and/or Elsevier’s API that were extracted using CADMUS (25) and converted to BioC-JSON files with Auto-CORPus (18).

We extracted the annotated entities from the results sections (IAO:0000318) and a list of each unique taxonomic identifier in these sections was extracted for each article. These lists were then aggregated per microbiome domain resulting in a list of identifiers and the number of times articles in that domain have reported them.

### Taxonomic tree

A combined set of all taxonomic identifiers found in any section of any document across the 14 domains was compiled and used to create a ‘reference’ taxonomic tree. The taxonomic identifiers were matched against the NCBI Taxonomy database (see ‘Data – NCBI Taxonomy’ section) and their parent IDs extracted. Any parent ID not yet in the list is then searched in the same database, until either it is part of the list or the parent ID’s rank is (super)kingdom (highest rank we include). Each entry in the reference taxonomic tree then gets the number of documents (count) it was found in as node weight.

For the domain specific trees, the structure of the reference tree is used with 0 counts for all nodes. Then the individual counts of microbes across domain-specific articles are added as node weights. Secondly, we traverse the tree, starting at the lowest taxonomic rank and add weighted counts to the parent node. The weighting takes place based on the number of child nodes the parent has in the complete taxonomy database, not only in the reference tree. As example, a genus with 10 children in the database and 3 children in the ‘reference’ will get an updated weight of their original node weight (counts) plus the counts of the 3 children divided by 10 children overall. If the genus has a weight of 5, and the 3 children have weights of 1, 0 and 3, respectively, the updated score of the genus is 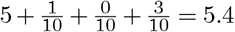.

Once the taxonomic tree is completely traversed and node weights updated, we then determine an empirical estimate of the null distribution of the node weights by resampling (nr=99,999 times, random state set to 2 before starting) using the reference tree weights (per taxonomic rank) as sampling probability for each non-zero node in the domain tree. I.e. for all nodes of rank ‘species’ in the domain tree we first sum the counts (‘sc’) and find the identifiers of these nodes (‘ni’). We then extract from the reference tree the counts for nodes with IDs contained in ‘ni’ as a vector, and normalise the vector (‘vp’) to a sum of 1 (probability). Random sampling with weights ‘vp’ is done ‘sc’ times for the rank, and these counts are added to the ‘i^th^’ resampled tree. This same process is done for all ranks. Any rank between any of the main ranks ((super)kingdom, phylum, class, order, family, genus, species) is sampled together with the higher rank (e.g. subgenus with genus). Then the counts in the resampled tree ‘i’ is aggregated as described above, and across all ‘nr’ resampled trees we count for each node the number of times (‘nt’) the aggregated count of the node is larger or equal to the domain tree’s aggregated count. This procedure allows us to estimate the proportion of times random sampling would result in higher counts than the actual value of the domain tree, and thus to calculate an empirical p-value for each node in the graph. The empirical P-value is defined as 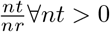 and 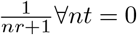. All P-values within each taxonomic rank are then adjusted for multiple testing using the Storey-Tibshirani False Discovery Rate (FDR, Q-value) given that they are independent at each rank but there is dependence between ranks. The Q-values are visualised on a *−log*10 scale, with significance indicated by Q*<*0.05 (or 1.30 on the *−log*10 scale). For any node in the graph, a significant Q-value indicates that this microbe is mentioned more than would be expected by random chance.

### Code and data availability

NCBI Taxonomy data is available from the [NCBI FTP server](https://ftp.ncbi.nlm.nih.gov/pub/taxonomy/new_taxdump/). The annotation pipeline code, code to train and inference NER, and code to inference NEN will be made available from [GitHub](https://github.com/omicsNLP/microbiomeNER). The fine-tuned models will be available from [HuggingFace](https://huggingface.co/). The OA (licences permitting redistribution) training set for NER will also be made available from [Zenodo](https://zenodo.com/) at this time together with the public part of the redistributable OA test set, with the private portion of the test set will be hosted on [CodaLab](https://codalab.org/). The list of identifiers in the reference tree and each domain tree will be made available through a separate [Zenodo](https://zenodo.com/) repository. Code to map results from a single study to an existing tree graph (reference or domain specific) will be made available from a separate [GitHub](https://github.com/) repository.

## Results

### Taxonomy dictionary and corpus creation

We curated a dictionary containing only the microbial (archaea, bacteria, fungi) entities from the NCBI Taxonomy database consisting of a total of 125,097 unique names mapping to 94,106 unique identifiers (Table 1), from (super)kingdom to species level. This dictionary was used as input to our annotation pipeline to automatically annotate 1,410 full-text articles for training and validation of deep learning models. A total of 90,150 annotations were made across the documents, with a median number of annotations between 51 and 61 for each of the five categories (Table 2). The majority of entities in the dictionary are fungi, however most annotations in the full-text documents are bacteria.

### Model Evaluation for Entity Recognition

We tested the models on a gold-standard test set of 288 documents that were annotated by two independent human researchers, with disagreements settled by a third arbiter. The inter-annotator agreement (across 20,486 annotations), represented by the F1-score, was 99.52% for NER. The pipeline without applying normalisation (i.e. exact matching of the recognised entity to an entry in the dictionary to filter out false positives) achieved a strict F1-score of 86.49% on the test set, and a relaxed F1 of 91.90% (see Table 3). Adding normalisation into the pipeline increases the performance to a strict F1 of 93.08% and relaxed F1 of 93.87%.

**Table 3.**
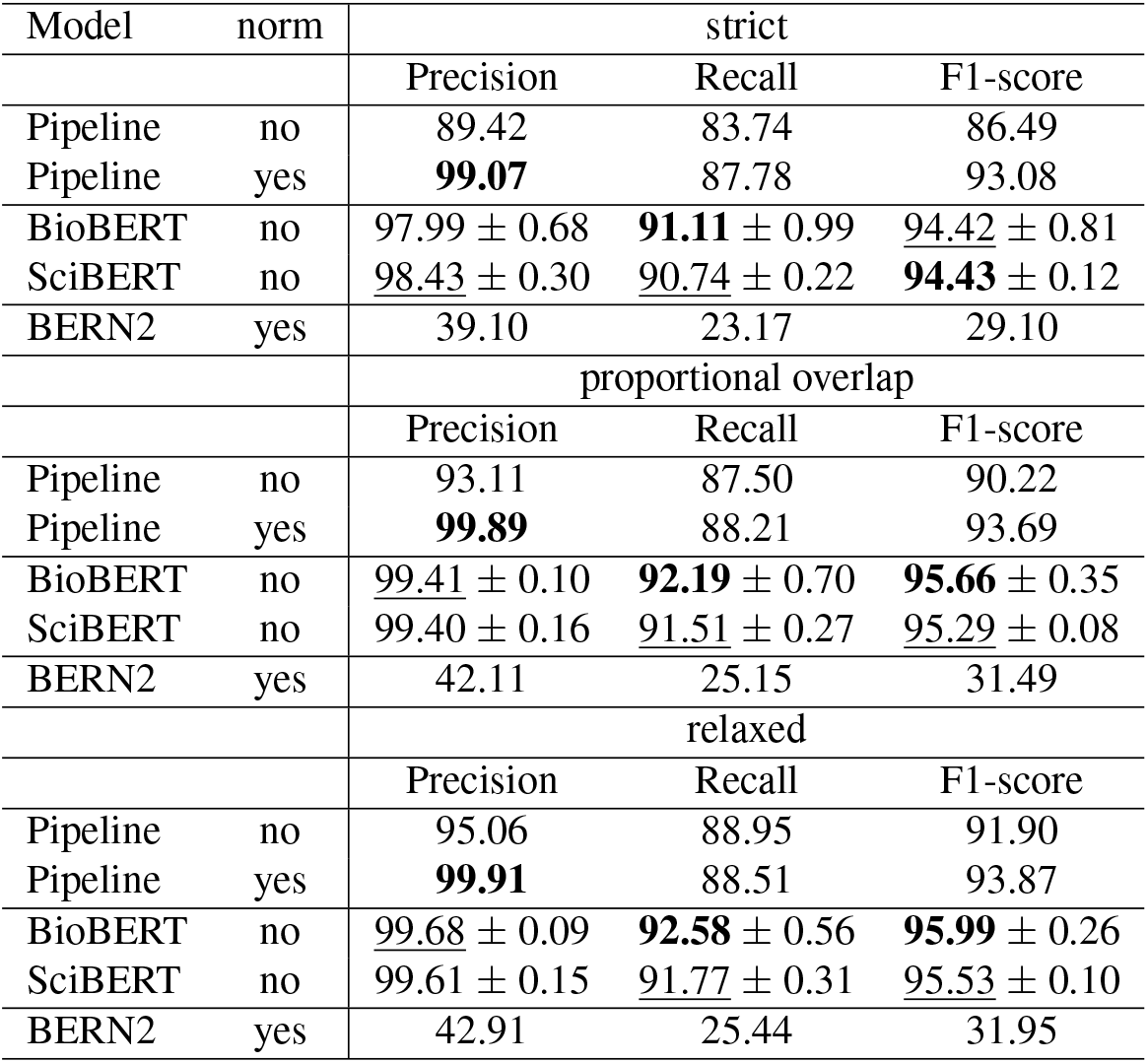
Performance metrics for different models (name and yes/no normalisation as part of entity recognition) evaluated on the manually annotated test set (n = 288 full text articles, total of 20,486 annotations) for entity recognition. Each metric (precision, recall, F1-score) expressed as percentage. Strict = indicates the complete span must match to be counted as true positive (TP), relaxed = at least one character must match between spans to be counted as TP, proportional overlap = overlap between two spans is calculated proportionally with number of matching tokens as TP, tokens missing as false negative, and unmatched tokens as false positives, normalised by the combined number of tokens length. Evaluation done excluding all entities that relate to the (super)kingdoms of bacteria, archaea and fungi as these are the most common found entities and including these would inflate numbers.

| Model | norm | strict |  |  |
| --- | --- | --- | --- | --- |
|  |  | Precision | Recall | F1-score |
| Pipeline | no | 89.42 | 83.74 | 86.49 |
| Pipeline | yes | **99.07** | 87.78 | 93.08 |
| BioBERT | no | 97.99 ± 0.68 | **91.11 ± 0.99** | 94.42 ± 0.81 |
| SciBERT | no | 98.43 ± 0.30 | 90.74 ± 0.22 | **94.43 ± 0.12** |
| BERN2 | yes | 39.10 | 23.17 | 29.10 |
| proportional overlap |  |  |  |  |
| Model | norm | Precision | Recall | F1-score |
|  |  | Precision | Recall | F1-score |
| Pipeline | no | 93.11 | 87.50 | 90.22 |
| Pipeline | yes | **99.89** | 88.21 | 93.69 |
| BioBERT | no | 99.41 ± 0.10 | **92.19 ± 0.70** | **95.66 ± 0.35** |
| SciBERT | no | 99.40 ± 0.16 | 91.51 ± 0.27 | 95.29 ± 0.08 |
| BERN2 | yes | 42.11 | 25.15 | 31.49 |
| relaxed |  |  |  |  |
| Model | norm | Precision | Recall | F1-score |
|  |  | Precision | Recall | F1-score |
| Pipeline | no | 95.06 | 88.95 | 91.90 |
| Pipeline | yes | **99.91** | 88.51 | 93.87 |
| BioBERT | no | 99.68 ± 0.09 | **92.58 ± 0.56** | **95.99 ± 0.26** |
| SciBERT | no | 99.61 ± 0.15 | 91.77 ± 0.31 | 95.53 ± 0.10 |
| BERN2 | yes | 42.91 | 25.44 | 31.95 |

Our two deep learning models were trained on labels generated by the pipeline with normalisation applied to the training and validation datasets (silver-standard). Our fine-tuned BioBERT and SciBERT models achieve strict F1-scores of 94.42 and 94.43, respectively, with relaxed F1-scores of 95.99 (BioBERT) and 95.53 (SciBERT). This gain in performance comes from an increase recall of the deep learning models, with precision closely matched to the pipeline with normalisation (99.91% vs 99.61-99.68%). The variance of the SciBERT model (across 9 models) is lower than that of the BioBERT model (Table 3), indicating it is more stable across different randomisations.

We evaluate the performance of the BERN2 model on the test set and compare it with our microbiome-specific models. Since not all species are microbial entities, and hence there can be many false positives, we evaluate BERN2 using the recall only. BERN2 achieves a relaxed recall of 25.44%, which is below our dictionary-based annotation pipeline (88.51-88.95%) and also not reaching the same level as our dedicated microbiome NER models (recall 91.77-92.58%).

### Model Evaluation for Entity Normalisation

Each of the annotations in the gold-standard test set (n=288 documents) was also given standard identifiers by two independent human researchers. The inter-annotator agreement across the 2,041 unique annotations was 88.31% (F1-score) for NEN. Disagreements were settled by a third arbiter as for NER to obtain a final test set with identifiers for each annotation. The pipeline achieved an accuracy for NEN of 89.64% across all 20,486 annotations, dropping to 69.28% when considering each unique annotation only once (see Table 4). Five deep learning models were evaluated for entity normalisation, 3 separate pre-trained models and our two fine-tuned models. These models have an overall accuracy for the top predicted identifier of 96.70-97.36% (Acc@1), increasing to 98.54-98.78% when a correct match is found in the top 5 predicted identifiers (Acc@5). For the unique accuracy, our fine-tuned models have an Acc@1 of 90.84-90.98% with the pre-trained models not far behind at 90.05-90.74%. At Acc@5, the pre-trained BioBERT model (95.00%) performs slightly better than our fine-tuned SciBERT (94.86%) and fine-tuned BioBERT (94.76%), with all models capable of accuracies *>*94%. Across all evaluations our fine-tuned SciBERT is top for the Acc@1 and second (after pre-trained BioBERT) for Acc@5. Hence, while the fine-tuned models show considerable improvement over pre-trained ones for NER, for NEN using BioSyn this does not appear to have any effect.

**Table 4.** Performance metrics (accuracy of top identifier (Acc@1) and top 5 identifiers (Acc@5) for different models evaluated on the manually annotated test set (n = 288 full text articles) for entity normalisation. Different model embeddings (existing and our fine-tuned models) were used to train BioSyn in hybrid mode.

| Model | fine-tuned | all |  | unique |  |
| --- | --- | --- | --- | --- | --- |
|  |  | Acc@1 | Acc@5 | Acc@1 | Acc@5 |
| Pipeline | N/A | 89.64 | N/A | 69.28 | N/A |
| BioBERT (3) | no | 97.13 | **98.78** | 90.74 | **95.00** |
| SciBERT (4) | no | 96.91 | 98.54 | 90.25 | 94.66 |
| SapBERT (24) | no | 96.70 | 98.68 | 90.05 | 94.02 |
| BioBERT | yes | 96.79 | 98.63 | 90.84 | 94.76 |
| SciBERT | yes | **97.36** | 98.70 | **90.98** | 94.86 |

### Using the NER and NEN models for literature review

The 6,927 full-text articles were annotated using the NER (fine-tuned BioBERT) and NEN (fine-tuned SciBERT embedding of the entity) pipelines, with the top candidate selected from the list of possible identifiers for each annotation. This resulted in a total of 182,064 entities in these documents, of which 6,014 were unique, and specifically 126,842 entities (4,878 unique) were found in the results sections. Resulting in a median number of reported unique entities in results sections of 30 (interquartile range 14-49), with a range of 0-578. Of the articles with at least one annotation identified in the results section text, 90% of articles have between 4 and 109 unique annotations mentioned.

Figure 1 displays the taxonomic trees for each of the 14 domains. Across all trees, most of the significant taxa (q*<*0.05) are either genera or species, followed by families and orders. The top 10 features per domain are given in Figure 2, with any feature that is significant in any domain and present in any domain’s top 10 visualised. This illustrates the similarity of certain domains that can also be appreciated from the taxonomic trees, but here from the view of individual microbes with *Lactobacillus, Streptococcus*, and *Prevotella* the most commonly reported taxa amongst those that are significantly overrepresented in at least one domain.

**Fig. 1.**
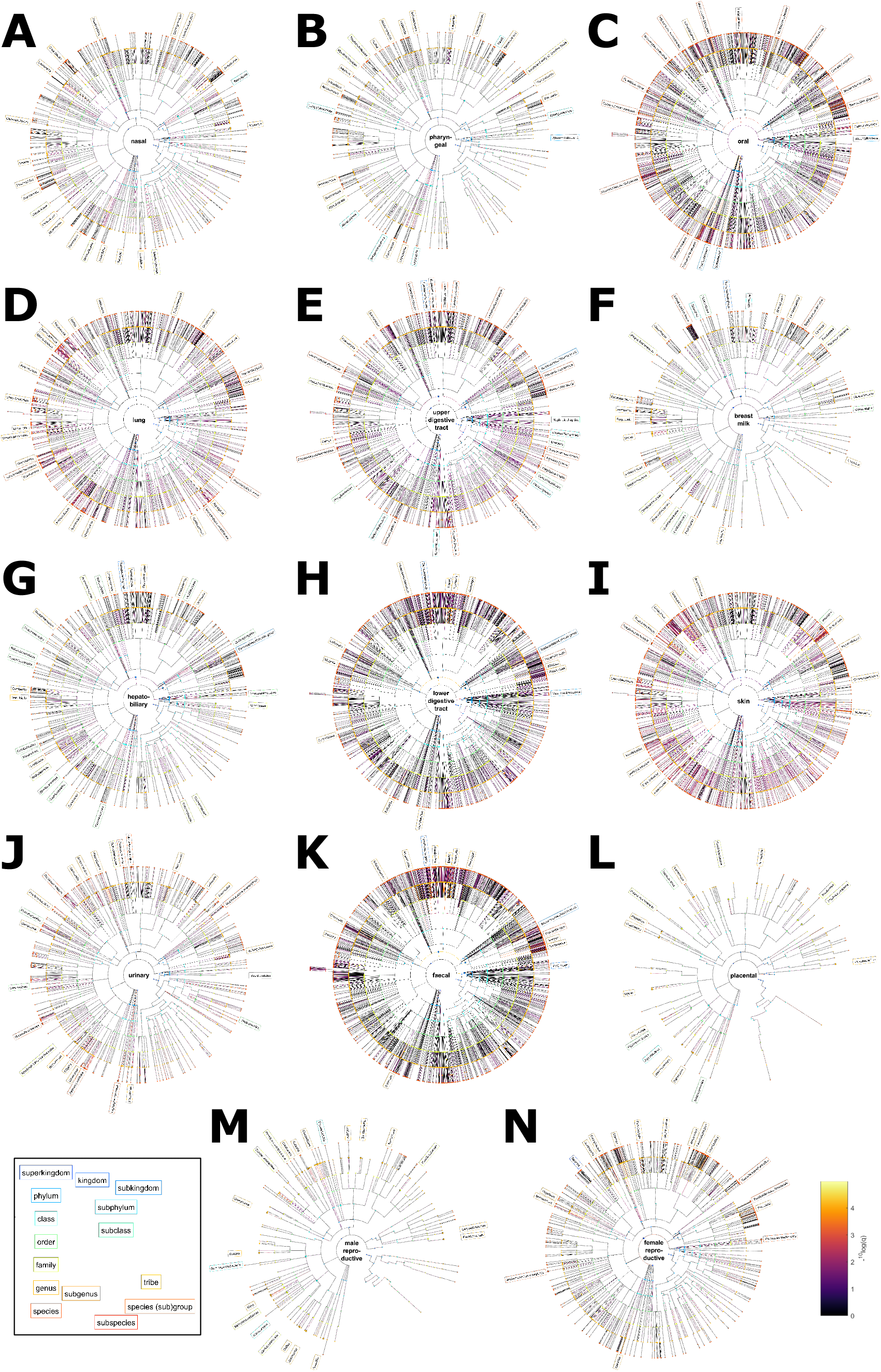
Taxonomic tree visualisation of domain-associated microbes. A) Nasal. B) Pharyngeal. C) Oral. D) Lung. E) Upper gastrointestinal. F) Breast milk. G) Hepatobiliary. H) Lower gastrointestinal. I) Skin. J) Urinary. K) Faecal. L) Placental. M) Male reproductive. N) Female reproductive microbiome. Individual high resolution figures for each domain can be found in the Supplementary Information. Colour of the nodes relate to the taxonomic rank (see legend). The colour of the edges is proportional to the −log10 of the q-value (higher meaning more significant). The top microbial entities associated with each domain are visualised around each graph and coloured based on the taxonomic rank.

**Fig. 2.**
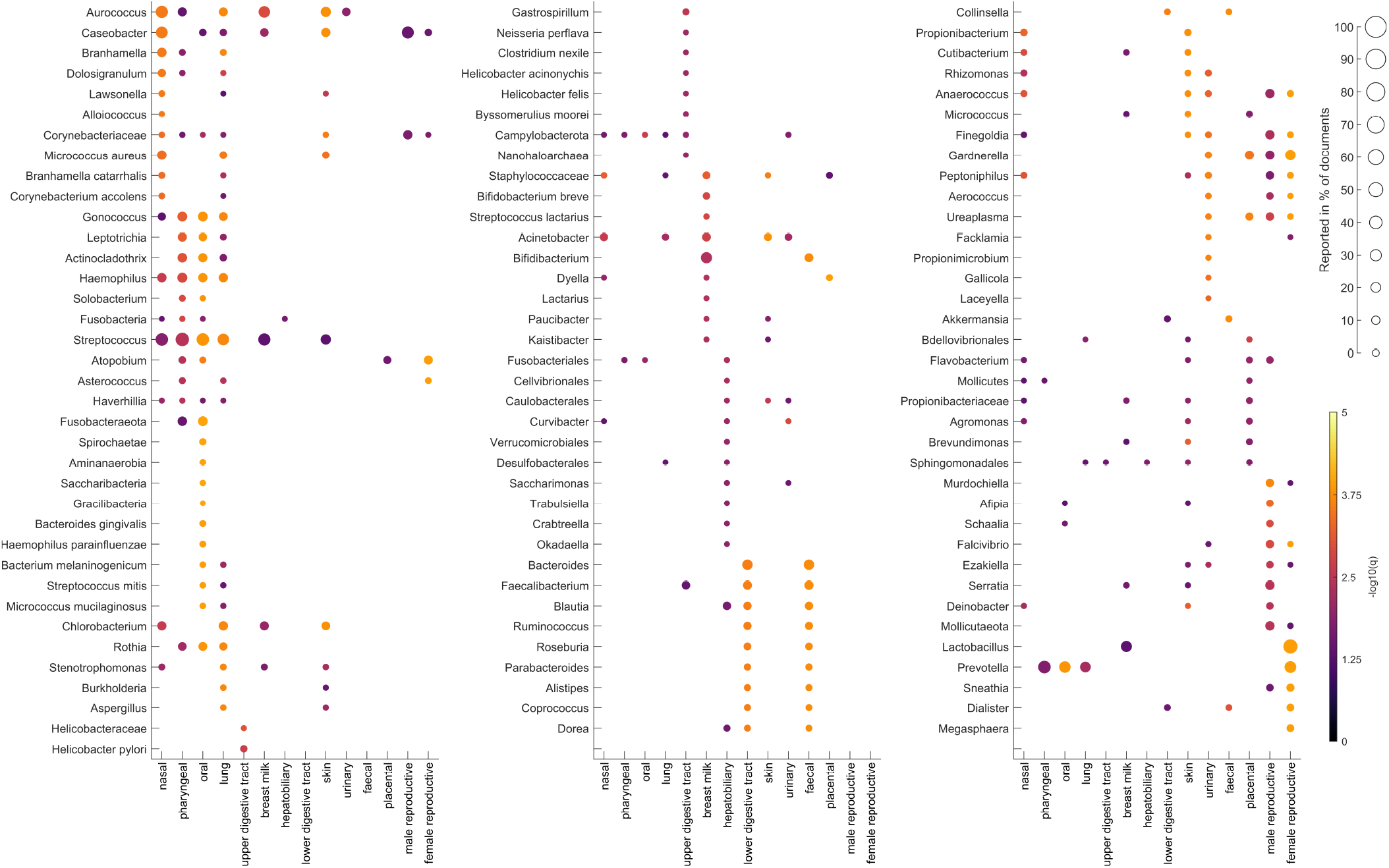
Heat map visualisation of the top 10 microbes for each of the 14 domains. Microbes with count *>* 1 and Q-value *≤* 0.05 are visualised. The size of the marker indicates the proportion of documents within the domain that report the microbe. The marker colour indicates the −log10 Q-value (higher meaning more significant).

### Common microbes

*Lactobacillus* was the most commonly reported taxon within a specific domain, with 73% of articles relating to the female reproductive (vagina, cervix, endometrium, but not urinary) microbiome reporting it in the results sections. It is significantly (Q=1.18×10^−4^) overrepresented in this domain compared to what would be expected at random. The only other domain in which it is significantly associated with is breast milk (human milk, colostrum, lactation) (Q=4.59×10^−2^), in 45% of articles. Within this genus *L. iners, L. crispatus, L. gasseri*, and *L. jensenii* (all at Q=1.57×10^−4^) were the most common species (26-46% of articles), with *L. johnsonii* (Q=2.08×10^−2^) and *L. psittaci* (Q=2.97×10^−4^) reported by 1-3% of studies on the female reproductive microbiome.

*Streptococcus* was found robustly associated (Q=1.43×10^−4^ to 4.98×10^−2^) with the oral (mouth, saliva), lung (lung, respiratory tract), pharyngeal, nasal (including nasopharynx), breast milk, and skin (skin, toe, foot, elbow fold, forehead) microbiomes (39-66% of articles). Individual *Streptococcal* species were reported in 0.3-7% of articles, with *S. mitis, S. salivarius, S. crista, S. vestibularis, S. peroris*, and *S. australis* reported by studies on the oral microbiome and at least one other domain (mostly lung, but also skin and upper digestive tract (oesophagus to duodenum) microbiomes).

*Prevotella* was most significantly reported in the female reproductive microbiome literature (Q=1.18×10^−4^, 51.6%), but also with oral, lung, and pharyngeal studies (Q=1.43×10^−4^ to 1.70×10^−2^, 42-59%). However, the species within the genus showed no overlap between these domains, with *P. bivia, P. amnii, P. timonensis, P. buccalis, P. disiens*, and *P. bergensis* (Q=1.57×10^−4^ to 5.66×10^−3^) reported in studies of the female reproductive microbiome, and *P. melaninogenica, P. pallens, P. oris, P. salivae*, and *P. jejuni* (Q=1.19×10^−4^ to 4.77×10^−2^) in common between oral and lung microbiomes (with no species significantly reported more often in relation to the pharyngeal microbiome).

Several other genera (e.g. *Neisseria, Haemophilus, Streptobacillus*) are in common between the nasal, pharyngeal, oral, and lung microbiomes, reflecting that these microbiomes are closely connected. Though there is some overlap with other domains for other taxa such as *Corynebacteriaceae* (skin, and male (testicle, semen, but not urinary) and female reproductive microbiomes) and *Campylobacterota* (upper digestive tract, urinary), with these taxa likely representing different lower rank taxa between these different domains. Overall, the most closely overlapped domains are faecal (faecal, stool) and lower digestive tract (jejunum to anus) microbiomes that have 462 different taxa associated significantly with at least one of these, of which 159 taxa (34%) are significantly associated with both. Despite that it appears the nasal, pharyngeal, oral, and lung microbiomes have a lot of overlap (Figure 2), in practice the ratio between overlapped taxa over the total is in the order of 9-13% (with oral and nasal having 4% overlap). Similar to that of the female and male reproductive systems (18/151, 12%), and the male reproductive (not including urinary) and urinary microbiomes (11%). The reproductive, urinary, and placental microbiomes have *Gardnerella* and *Ureaplasma* in common, with the male and female reproductive system and urinary microbiomes sharing other common taxa with skin including *Anaerococcus, Finegoldia*, and *Peptoniphilus*.

### Domain specific microbes

However, from the same data, certain taxa are only found overrepresented in a single domain. For example, a number of taxa are significant only for the oral microbiome such as the phyla *Spirochaetota, Synergistota, Saccharimonadota*, and *Altimarinota* (all Q=1.08×10^−4^). These were recognised in the text with different synonyms, e.g. for *Spirochaetota* (txid:203691) only one article used this nomenclature. The majority used *Spirochaetes* (n=132), with *Spirochaetota* (n=15), *Spirochaetae* (n=11), and *Spirochaeota* (n=2) used more often than that current accepted name.

The upper gastrointestinal tract also had considerable taxa that were found to be significantly associated only with this domain including *Helicobacteriaea, Helicobacter, H. pylori* (with 4 articles using synonym *Camphylobacter pylori*), *H. acinonychis, H. felis*, and *H. hominis* (Q=9.68×10^−4^ to 1.22×10^−2^), as well as *Neisseria perflava, Tyzzerella nexilis, Dictyonema moorei*, and *Nanohaloarchaea*.

Several *Bifidobacteria* were significantly reported more in the breast milk microbiome, such as *B. breve, B. choerinum, B. biavatii*, and *B. saguini* (Q=1.46×10^−3^ to 4.16×10^−2^). Also, significantly higher in only breast milk were *Streptococcus lactarius*, the fungus *Lactarius*, and *Plesiomonas*.

Finally, the hepatobiliary (liver, pancreas, bile duct, gall bladder) microbiome contains several significantly associated taxa at the order, *Cellvibrionales, Verrucomicrobiales*, and *Anaeroplasmatales* (Q=5.12×10^−3^ to 1.64×10^−2^), and genus levels, *Trabulsiella, Crabtreella, Okadaella*, and *Pseudochrobactrum* (Q=1.23×10^−2^ to 1.58×10^−2^). A complete table of all significantly associated microbes (common and unique) can be found in the Supplementary Materials.

## Discussion

This work is the first to create microbiome-specific corpora and algorithms for text mining research. Upon peer review, we will share the annotated Open Access documents that have licenses permitting redistribution in two ways. First, we share the training set with labels from our pipeline freely. Second, our gold-standard test set is split into a private (50 documents) and public (remaining documents) allocation. The private set is hosted on CodaLab with unannotated and normalised versions of the documents available for others, this is set up to allow benchmarking against these documents and to prevent data leakage of test data into models. The public set can be used to locally evaluate models, however with the risk that these data can be ingested into the training of LLMs (data leakage). Like the training set, we also share all documents with annotations and linked identifiers that made up our dataset for literature review, these documents were annotated by our DL algorithms (and not the pipeline). Our corpora are larger than most bioNLP corpora (26), and compared with species-only corpora (27, 28) our training corpus contains 20× more annotations. The number of full-text documents contained in our corpora also exceeds that of the full-text (27) and abstract-only (28) species corpora. The results sections alone from the literature review corpus contain 182,064 annotations (6,014 unique), thus providing a valuable resource of additional training data for training NER and NEN algorithms for recognising microbial entities.

The training corpus was annotated using the pipeline and used as input to train DL algorithms as an alternative to dictionary searching combined with rule-based annotation. The pipeline algorithm itself has been used separately as a first-pass approach to annotate microbiota followed by human review for a meta-analysis of microbiome research on malnutrition in African populations (29). We have previously demonstrated how ‘silver’ annotated training data can achieve accurate NER algorithms for domains that have no training data available such as metabolites (30) and enzymes (20). Here we showed a performance gain of over 2% for NER of the DL algorithms (BioBERT achieving an F1-score of 0.96), and over 30% for NEN (SciBERT achieving an accuracy of 91%). While the DL models have slightly lower precision than the pipeline (−0.3%) for NER, this is negligible compared with the gain in recall and speed. This indicates that the DL models learn how to identify entities in the sentence rather than only identifying entities present in the training set. The pipeline takes on average 202 seconds compared with the DL models taking only 7 seconds per document. A challenging aspect of our test set is that 7.73% of annotations in the test set can only be found here and are not contained in the training set. The higher recall of the DL models demonstrated that these models are able to recognise additional entities not found using exact matching, including some spelling errors in names that were correctly resolved to the right taxonomic identifier with the NEN models. This model can also be used in conjunction with others (mixture of experts) for annotation of multiple categories of entities. For example, one of the limitations of our prior work on metabolite NER(30) demonstrated that a small proportion of microbiota are falsely recognised as metabolites. This likely occurs due to the similarity of certain tokens such as for example ‘butyr’ in the metabolite butyrate and the microbial genus *Butyrococcus*.

Filtering the data for any significant taxa (Q*<*0.05) reported at least by 2 independent studies showed that a coverage of at least 50% of all these significant taxa can be obtained with only 5 studies for most domains. The exceptions were faecal (with 5 studies combined mentioning 46% of the 300 significant taxa), skin (38%, 298 significant taxa), and upper digestive tract (38%, 114 significant taxa), with 6, 9 and 8 articles needed to have a coverage of at least 50% for each of these respectively. While this is logically related to the number of unique taxa reported per domain, others with similar numbers (e.g. oral with 292 significant taxa) can have a coverage of over 50% with fewer articles. This strategy can be used to order relevant articles for the literature review stage of a new study. The way we envision this process can take two forms. The first is analogous to how it was done here, by obtaining machine-readable formats for all relevant articles through a PubMed search and obtaining XML versions of these full-text documents (25), conversion of such documents to the machine-readable BioC-standard (18), followed by annotation using the NER and NEN models contributed here. From these the reported taxa from results sections can be extracted and summarised, and a document ranking achieved based on the unique number of taxa each article contributes (the background set of microbes reported in any domain, or in specific domains can both be used for this purpose). For example, for the vaginal microbiome the combination of 3 studies (31– 33) describes 51% of the microbes that we find significantly reported more often in this domain compared to others, this number which goes up to 83% with 10 articles (3.6% of the total of 281 articles in our corpus from this domain). This approach can lead to identifying targets to test in new data to reproduce earlier results, or to otherwise generate hypotheses. The second approach starts the same to collect all relevant articles (with appropriate search terms to filter more relevant articles at the first stage), however rather than finding taxa that are reported more often than would be expected by random chance instead the results from a new study can be compared against the literature to find the relevant articles to inspect further. While there will not be a BioC-version of a manuscript in preparation, this is not needed for the algorithms to work. I.e. if the significant microbes are passed as text file to the NEN it will output the taxonomic identifiers, and then based on the entire body of relevant literature a search can be conducted to find a set of articles that describe as much from the findings as possible. At the same time, this will also implicitly pinpoint any novel findings in a new study. For example, taking the most recent article in our corpus on the breast milk microbiome (34), we can establish that different combinations of 4-5 articles from our corpus report the 22 unique taxa reported in 34 (published in December 2023), thus any of these combinations could form a logical first step for literature review. For example, of a set of 4 (35–38) (April 2016 to May 2023), 2 of these were cited and of another (37) an article by the same authors was cited. Only one article published in the same year (May 2023) was not cited. Another combination that fully covers all reported microbes contains 5 articles (35, 36, 38–40), of which the same two were cited and 38 was not, but the other two were not cited however articles by the same authors were cited (likely covering similar cohorts). Using this approach to shortlist articles for review is dependent on adequately defining the relevant literature (human effort) with the steps after it automated. The automation can save researchers time with identifying which articles to read first by generating a list of priorities. With domains that have hundreds (only the breast milk, male reproductive, and placental microbiomes had under 100 articles) or thousands (the faecal microbiome portion of our corpus contains over 2,000 documents) of articles, this approach could save time for researchers when conducting their literature review, as noted in previous work (29).

## Limitations

One limitation of our methods are that they were not trained to recognise viruses, but only trained on archaea, bacteria, and fungi. This was recognised by 29 whom annotated the viruses manually. However, as viral genomes are typically much smaller and lacking a universal marker gene (e.g. 16S or 18S rRNA, ITS) they are not commonly captured by the bioinformatic pipelines used to compare sequences with libraries. With the released corpora and pipeline, viruses can be added to the pipeline for these documents to be re-annotated. We share our code and models (following peer review) to retrain or fine-tune the DL models thus these can be updated to include viruses. However, since our test set does not contain viruses, a new test set would need to be created to evaluate the accuracy of the updated models. Here, we used BERT-based architectures as opposed to LLMs. LLMs have been shown to fail to generalise to out-of-domain tasks such as entity recognition, with no LLM coming close to BERT-based models for NER (8–10) or NEN (9). In the initial model development stage we explored different open-weight, locally-deployable LLMs alongside the BERT models, however have found them to lag behind as literature suggested for NER and did not pursue this further. The BERT-based models are capable of annotating entire documents in seconds, however this does require such models to be loaded into memory beforehand. Hence, annotating a single document will take considerably longer than annotating many documents, since the model loading time cannot be amortised across multiple inputs. Likewise, loading the NEN model also costs time but inference is fast.

Utilising these models for literature summarisation has several aspects that have room for improvement. First, it is dependent on defining a set of articles relevant to a new study, this human task would ideally be done in a systematic manner to determine appropriate inclusion and exclusion terms. The search queries used here can be adapted by adding additional filters such as e.g. filtering faecal microbiome studies for specific diseases of interest. Our models here focus on first recognising the entities and second assigning identifiers to them. They do not perform relation extraction, thus do not provide any context of how the identified microbe is related to the study outcome (e.g. is it higher/lower compared to controls). In other contexts, LLMs have shown capabilities to reliably extract relations (8, 9) thus these could be explored should users want to automate this step as well in favour of human review.

## Conclusions

In summary, this work contributed a novel dictionary- and rule-based automatic pipeline for microbiome NER and NEN with almost perfect precision. The best performing DL model was BioBERT with a state-of-the-art performance (F1-score*>*0.95) for microbiome NER that considerably speeds up the annotation process. We have shown the application of these models for assisting with literature review and capability to meta-analyse the existing literature to determine taxa that are significantly over-reported in certain domains. The code for automatic annotation, model training and inference, visualisation, and all data will be made available following publication.

## AUTHOR CONTRIBUTIONS

Conceptualisation: JP; Methodology: DP, AL, and JP; Software: DP, AL, AV, NFM, MW, and JP; Validation: DP, AL, AV, and JP; Formal analysis: DP, AL, AV, NFM, and JP; Investigation: DP, AL, AV, NFM, and JP; Resources: TB and JP; Data Curation: DP, AL, NFM, MM, and JP; Writing - Original Draft: JP and AL; Writing - Review & Editing: DP, AL, AV, MM, TB, and JP; Visualisation: JP; Supervision: TB and JP; Project administration: JP; Funding acquisition: TB and JP. All authors have read and agreed to the published version of the manuscript.

## Funding

A.V. is supported by a UK Research and Innovation (UKRI) Centre for Doctoral Training in AI for Healthcare PhD studentship (EP/S023283/1 (EPSRC)). This research was funded by Health Data Research (HDR) UK and the Medical Research Council (MRC) via an UKRI Innovation Fellowship to T.B. (MR/S003703/1) and a Rutherford Fund Fellowship to J.M.P. (MR/S004033/1). T.B., J.M.P. and A.L. are additionally supported by the Horizon Europe project CoDiet. The CoDiet project is funded by the European Union under Horizon Europe grant number 101084642. UK participants in Horizon Europe Project CoDiet are supported by UKRI grant numbers 10060437 (Imperial College London) and 10102628 (University of Nottingham). J.M.P. and M.N.M. are supported by the MRC project GI-tools (MR/V012452/1). The authors declare no conflict of interest. The funders had no role in the design of the study; in the collection, analyses, or interpretation of data; in the writing of the manuscript, or in the decision to publish the results.

## Notes

### Competing Interest Statement

The authors have declared no competing interest.

### Summary of Updates

Updated GitHub link in the abstract which referred to an old repo.

