## Supplementary Information for "Microbial Named Entity Recognition and Normalisation for AI-assisted Literature Review and Meta-Analysis"

### Supplementary Figures

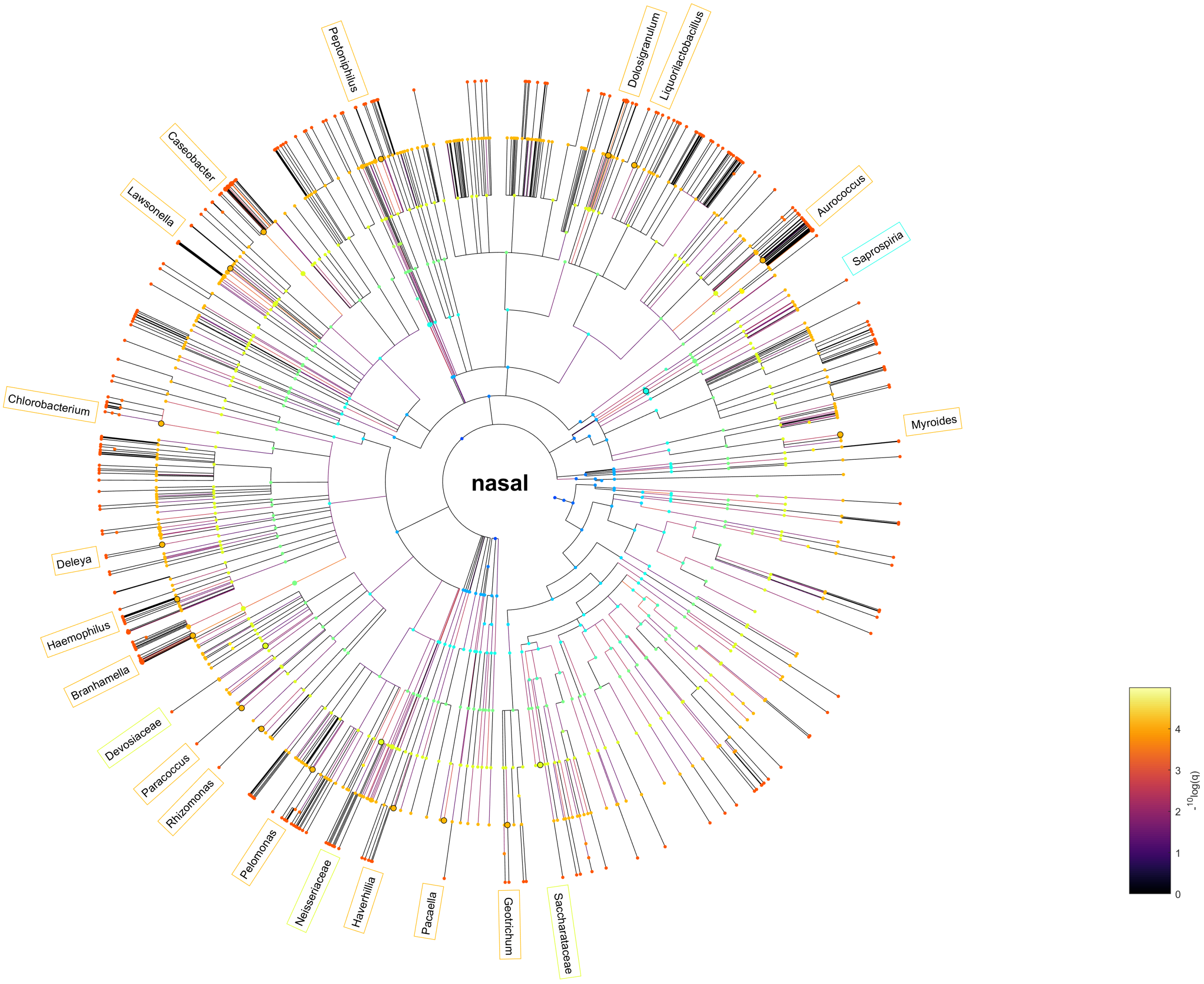


**Fig. 1.** Taxonomic tree visualisation of the nasal (including nasopharynx) microbiota. Colour of the nodes relate to the taxonomic rank. The colour of the edges is proportional to the -log10 of the q-value (higher meaning more significant). The top microbial entities associated with the domain are visualised around the graph and coloured based on the taxonomic rank.

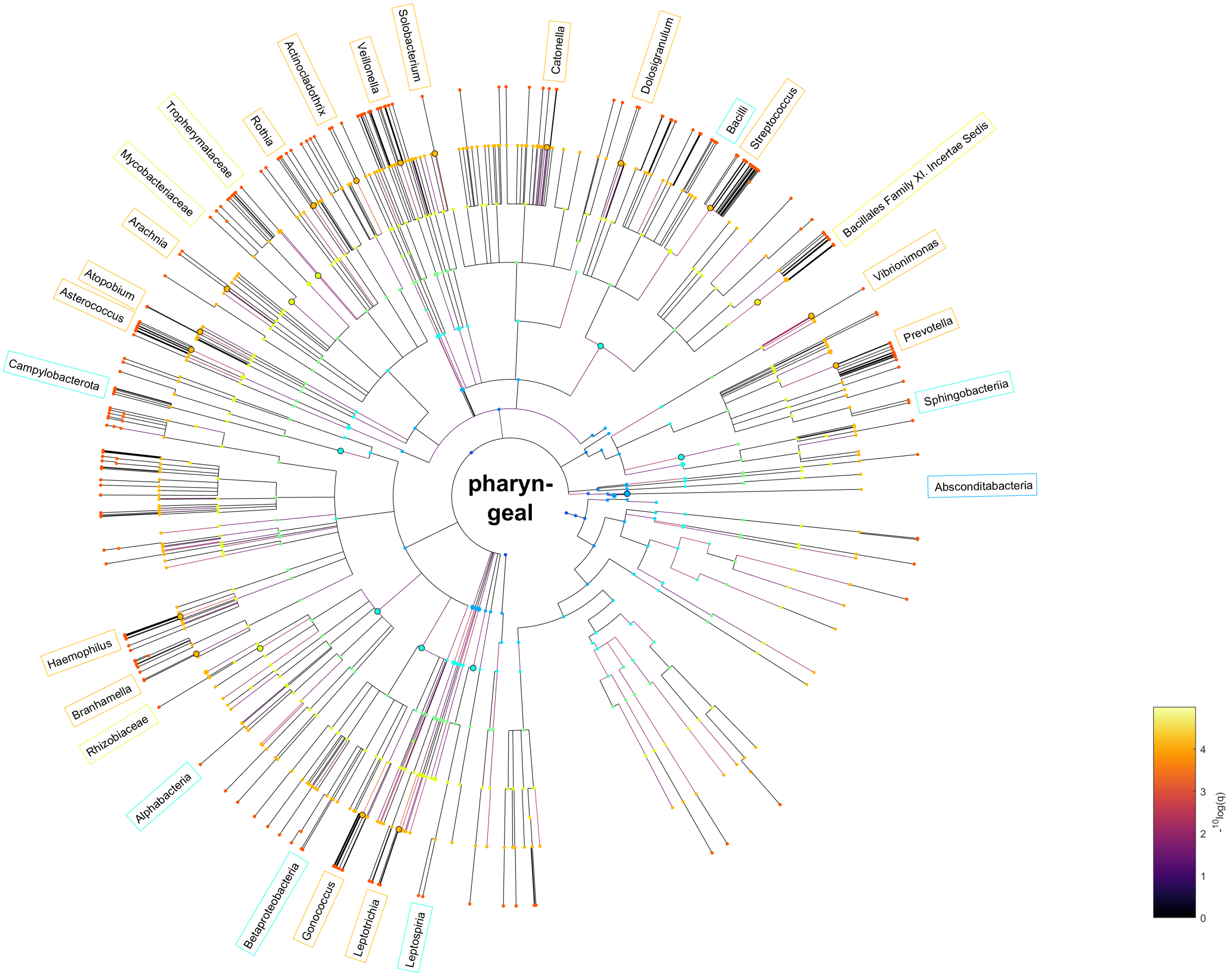


**Fig. 2.** Taxonomic tree visualisation of the pharyngeal microbiota. Colour of the nodes relate to the taxonomic rank. The colour of the edges is proportional to the -log10 of the q-value (higher meaning more significant). The top microbial entities associated with the domain are visualised around the graph and coloured based on the taxonomic rank.

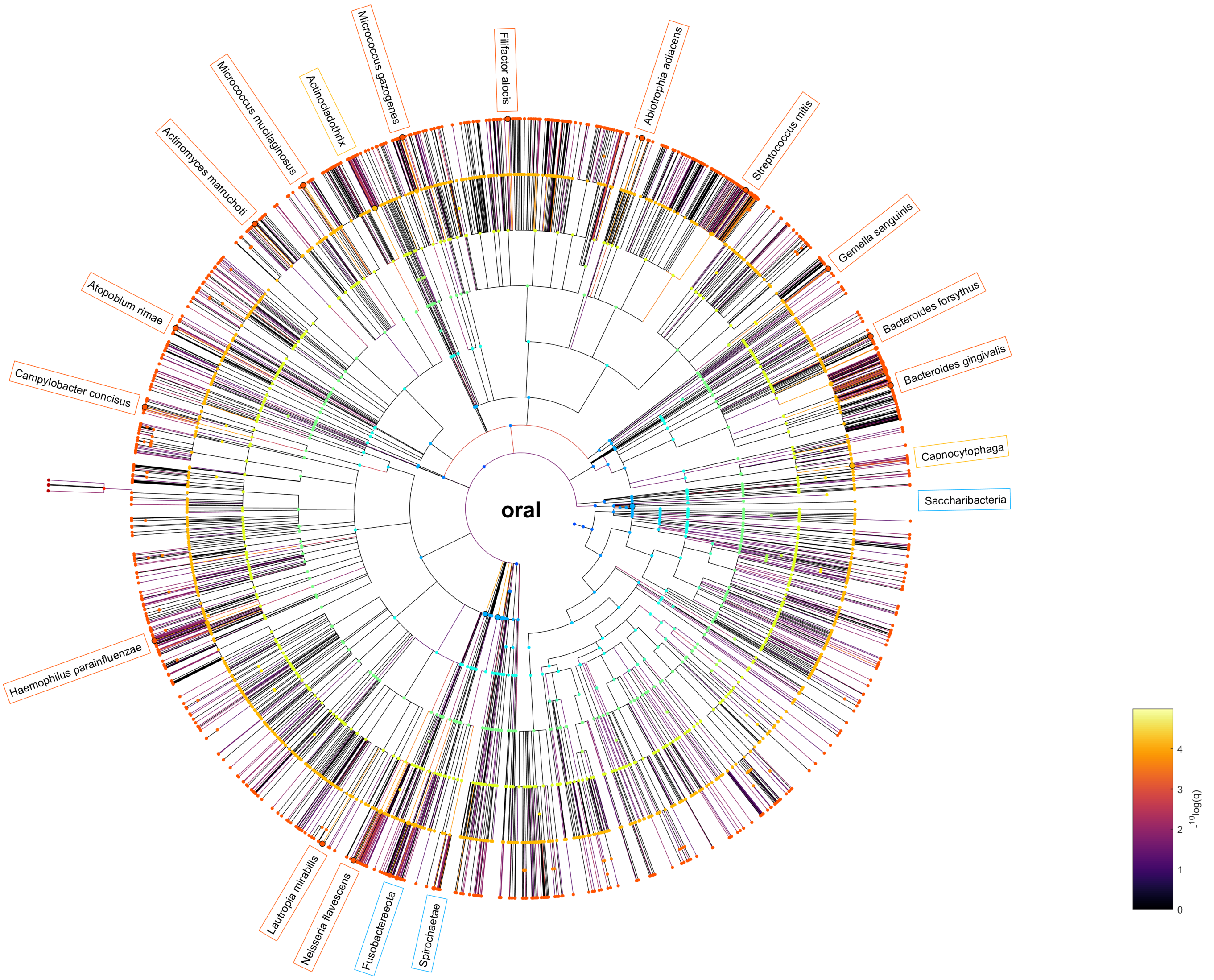


**Fig. 3.** Taxonomic tree visualisation of the oral (mouth, saliva) microbiota. Colour of the nodes relate to the taxonomic rank. The colour of the edges is proportional to the -log10 of the q-value (higher meaning more significant). The top microbial entities associated with the domain are visualised around the graph and coloured based on the taxonomic rank.

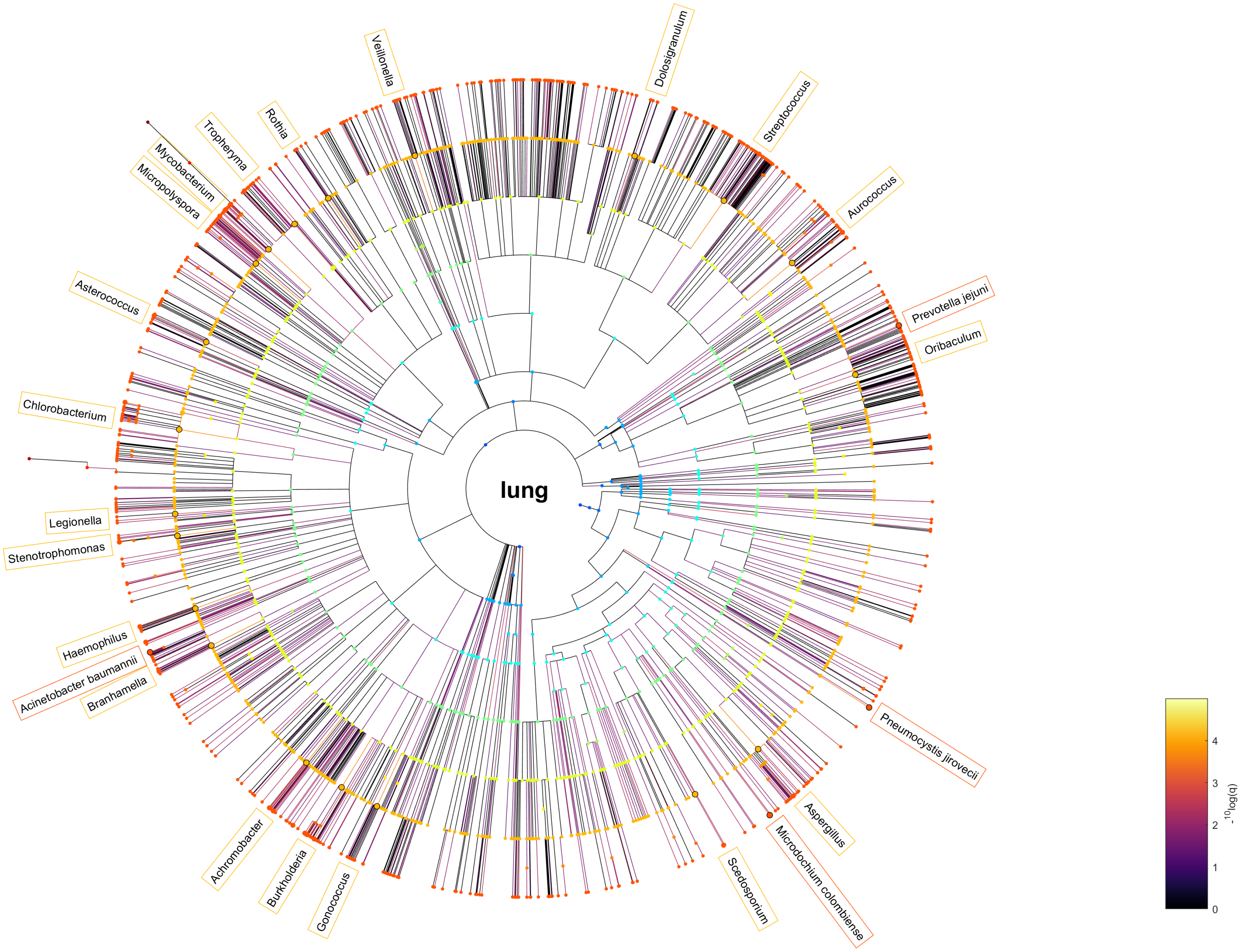


**Fig. 4.** Taxonomic tree visualisation of the lung (including respiratory tract) microbiota. Colour of the nodes relate to the taxonomic rank. The colour of the edges is proportional to the -log10 of the q-value (higher meaning more significant). The top microbial entities associated with the domain are visualised around the graph and coloured based on the taxonomic rank.

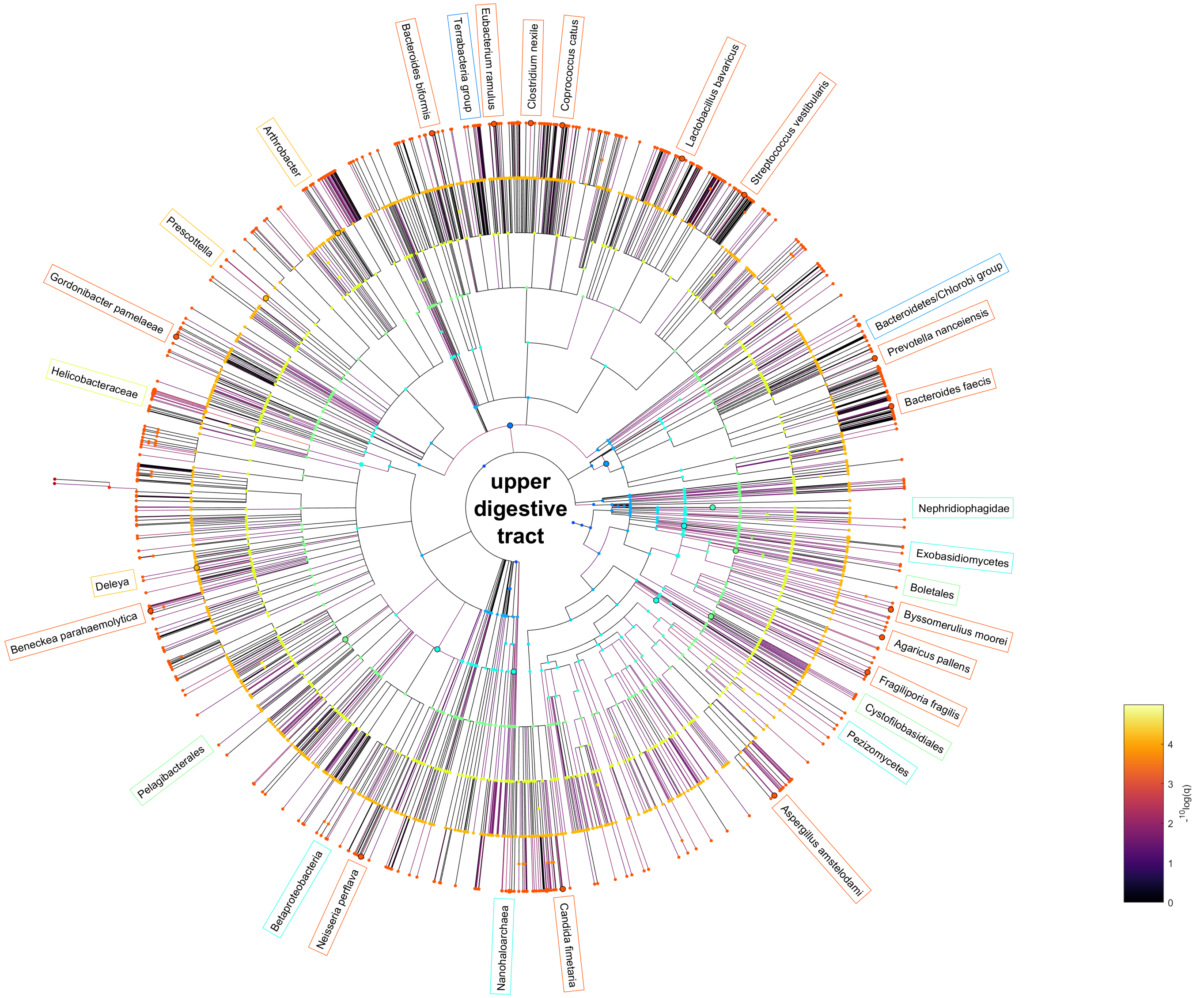


**Fig. 5.** Taxonomic tree visualisation of the upper gastrointestinal (oesophagus to duodenum) microbiota. Colour of the nodes relate to the taxonomic rank. The colour of the edges is proportional to the -log10 of the q-value (higher meaning more significant). The top microbial entities associated with the domain are visualised around the graph and coloured based on the taxonomic rank.

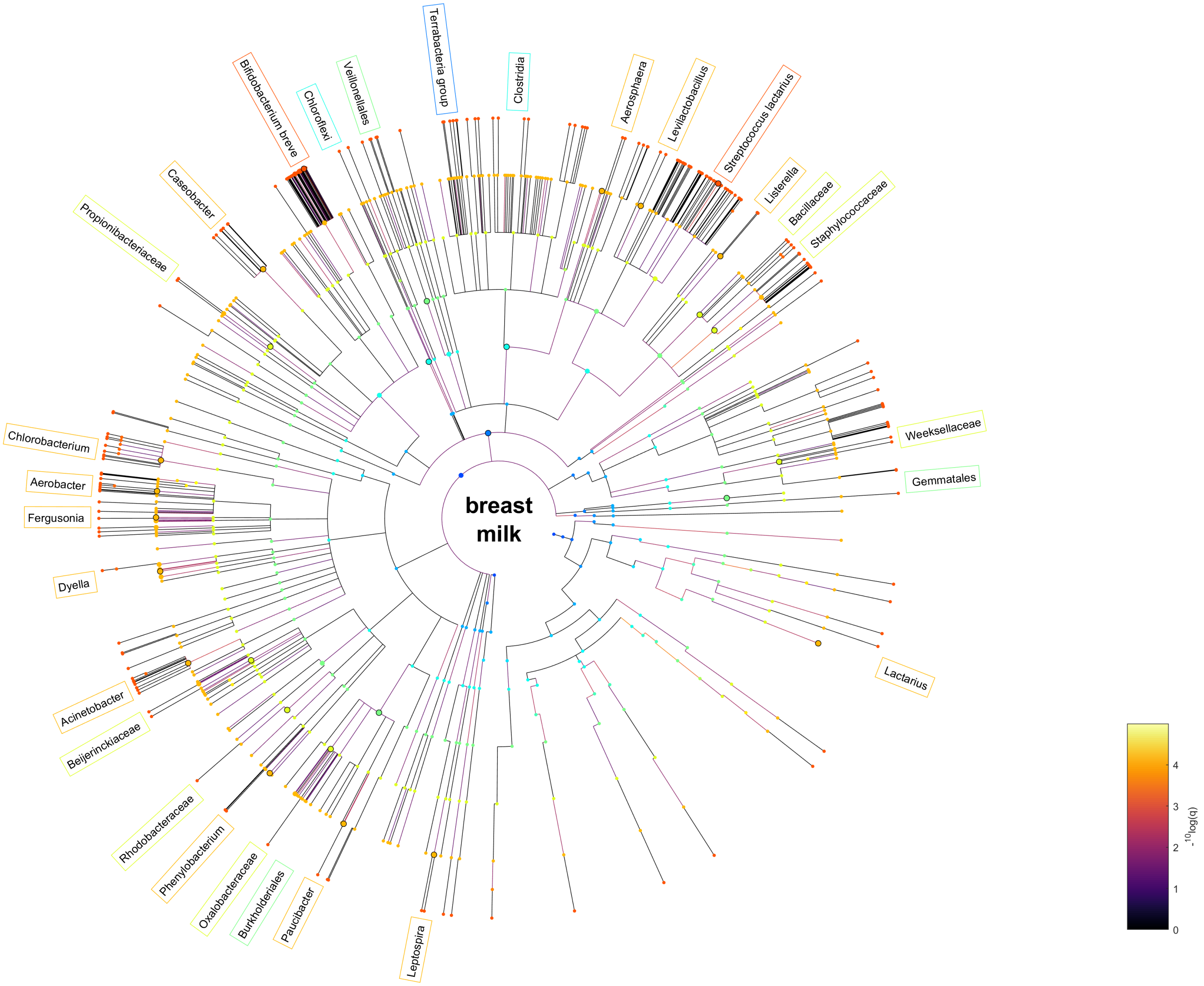


**Fig. 6.** Taxonomic tree visualisation of the breast milk (human milk, colostrum, lactation) microbiota. Colour of the nodes relate to the taxonomic rank. The colour of the edges is proportional to the -log10 of the q-value (higher meaning more significant). The top microbial entities associated with the domain are visualised around the graph and coloured based on the taxonomic rank.

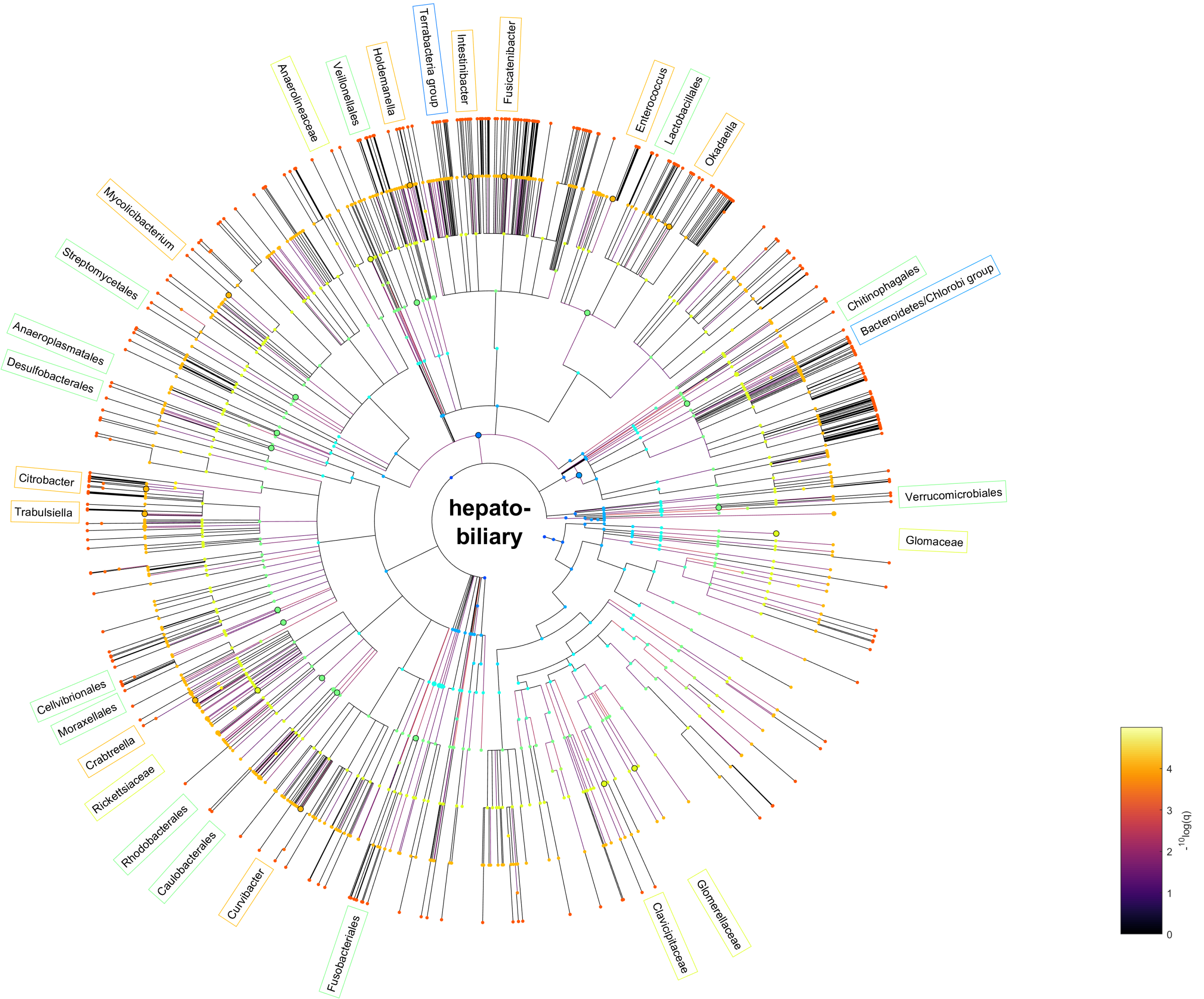


**Fig. 7.** Taxonomic tree visualisation of the hepatobiliary (liver, pancreas, bile duct, gall bladder) microbiota. Colour of the nodes relate to the taxonomic rank. The colour of the edges is proportional to the -log10 of the q-value (higher meaning more significant). The top microbial entities associated with the domain are visualised around the graph and coloured based on the taxonomic rank.

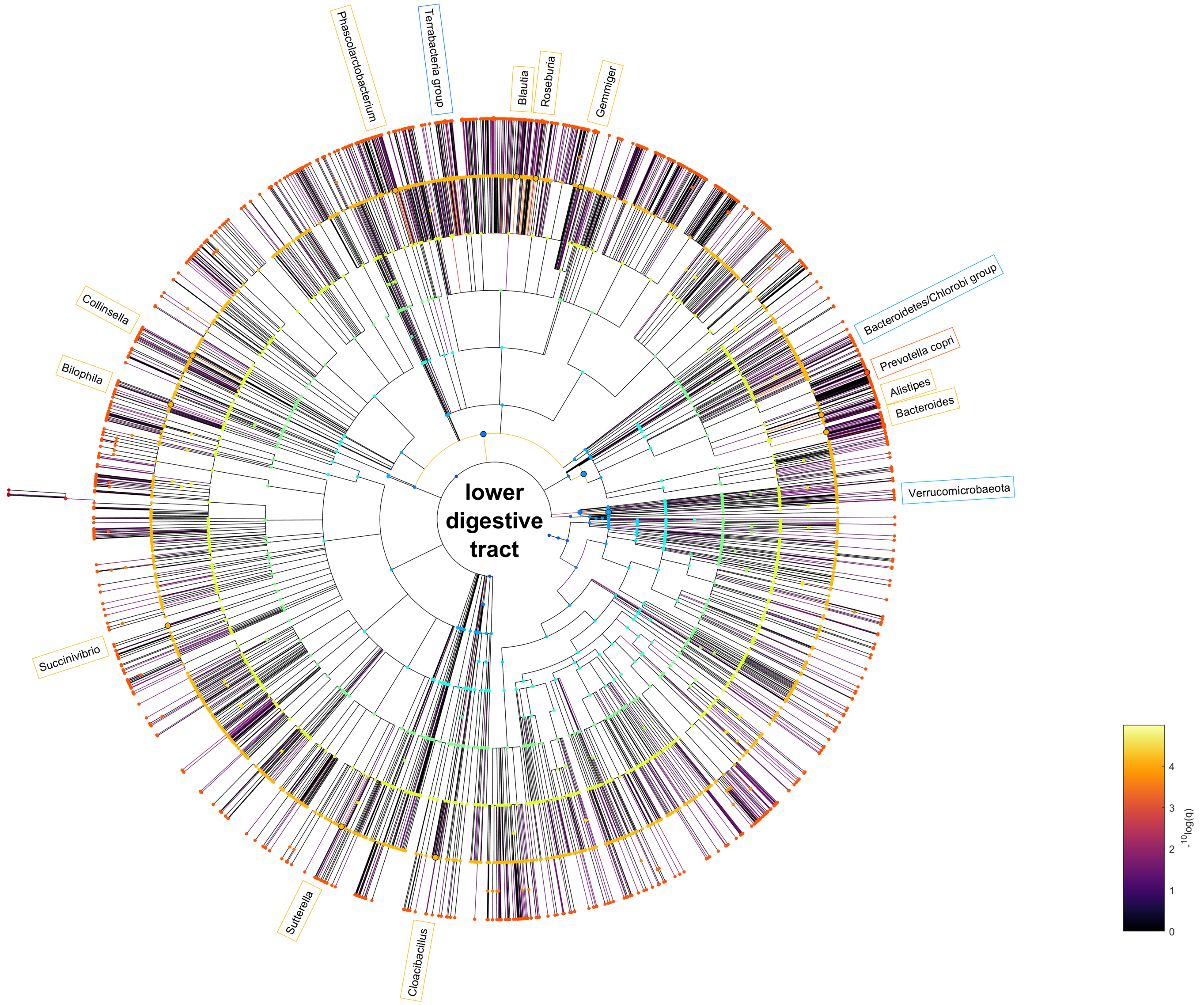


**Fig. 8.** Taxonomic tree visualisation of the lower gastrointestinal (jejunum to anus) microbiota. Colour of the nodes relate to the taxonomic rank. The colour of the edges is proportional to the -log10 of the q-value (higher meaning more significant). The top microbial entities associated with the domain are visualised around the graph and coloured based on the taxonomic rank.

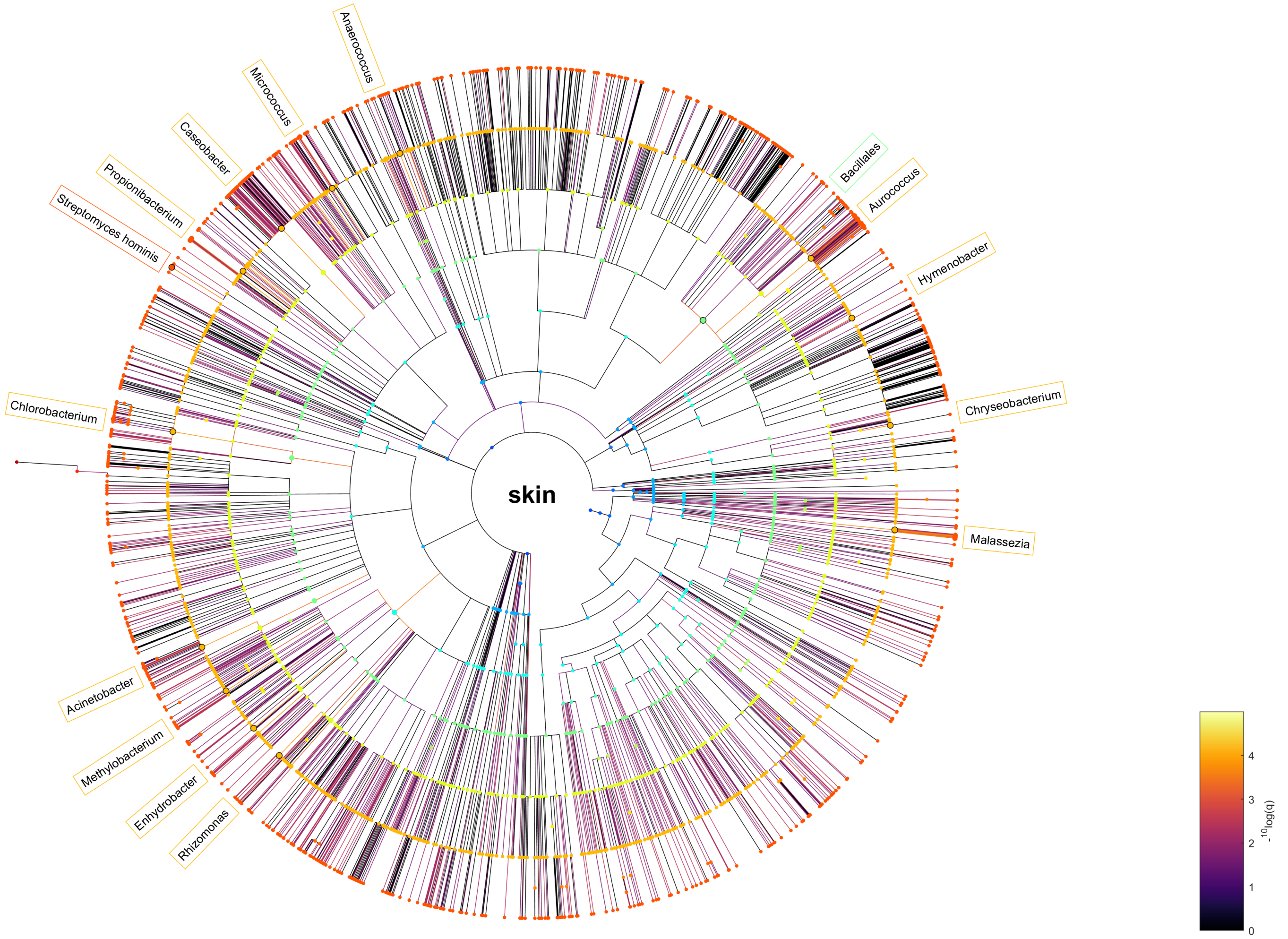


**Fig. 9.** Taxonomic tree visualisation of the skin (skin, toe, foot, elbow fold, forehead) microbiota. Colour of the nodes relate to the taxonomic rank. The colour of the edges is proportional to the -log10 of the q-value (higher meaning more significant). The top microbial entities associated with the domain are visualised around the graph and coloured based on the taxonomic rank.

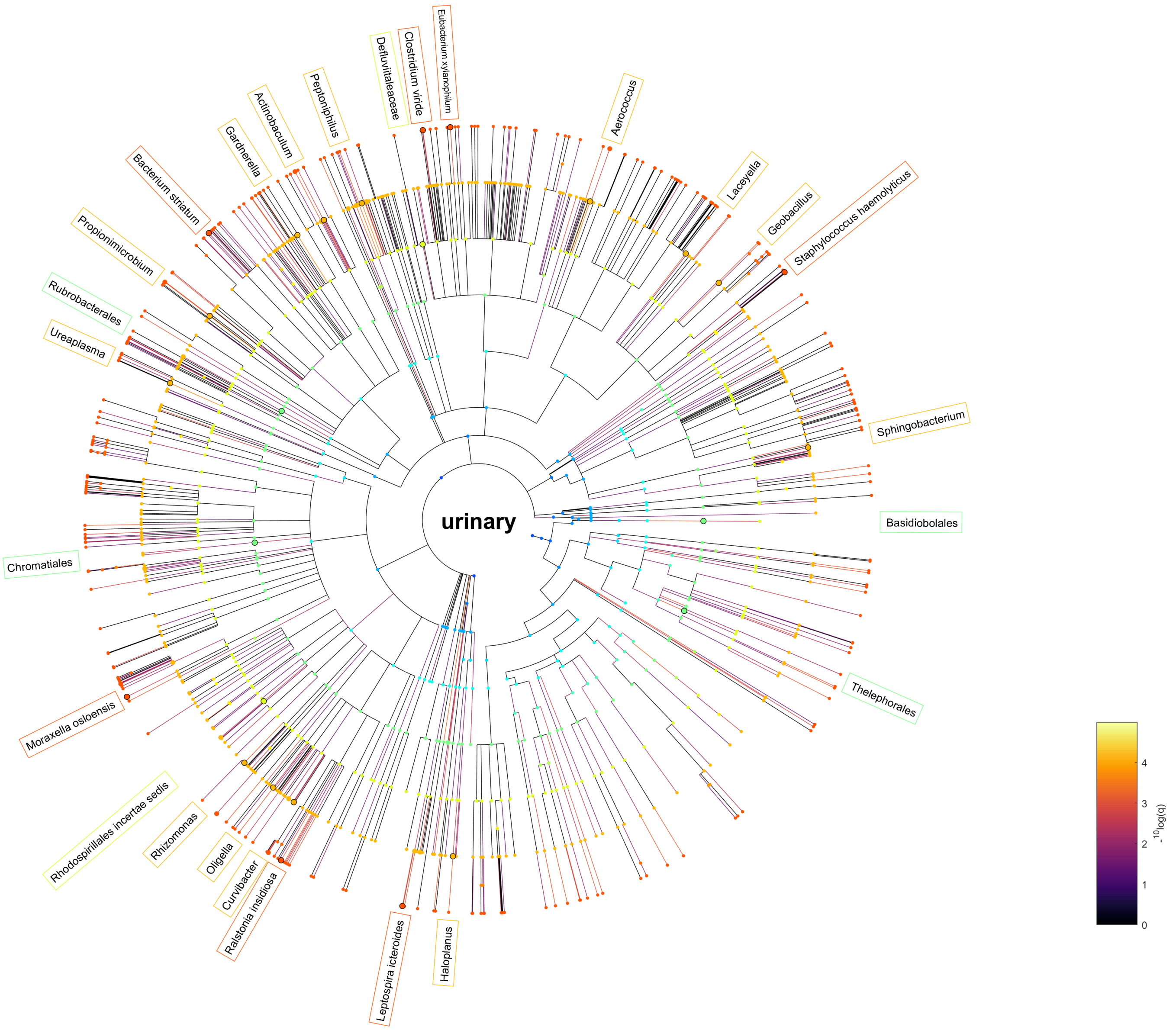


**Fig. 10.** Taxonomic tree visualisation of the urinary microbiota. Colour of the nodes relate to the taxonomic rank. The colour of the edges is proportional to the -log10 of the q-value (higher meaning more significant). The top microbial entities associated with the domain are visualised around the graph and coloured based on the taxonomic rank.

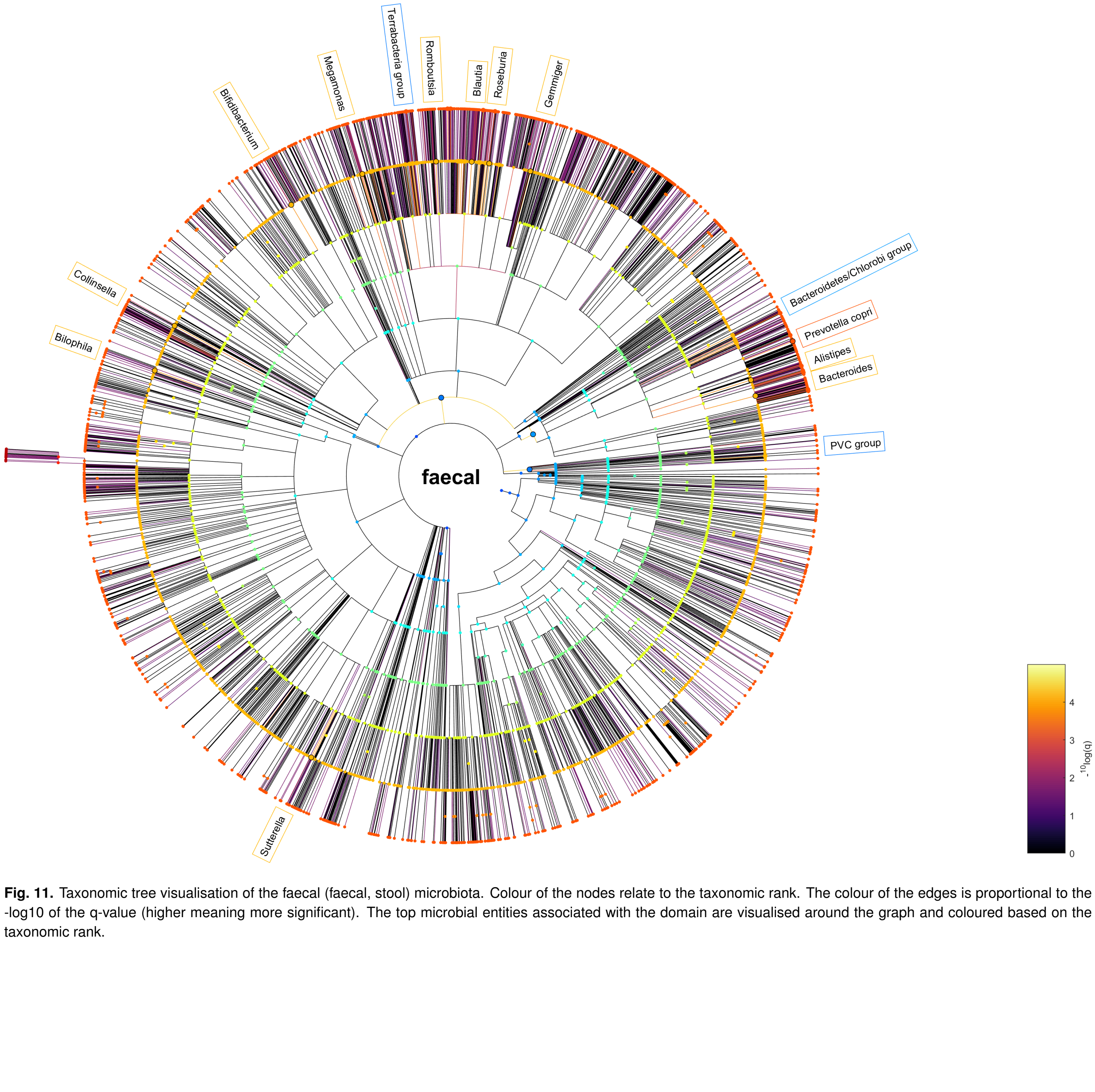


**Fig. 11.** Taxonomic tree visualisation of the faecal (faecal, stool) microbiota. Colour of the nodes relate to the taxonomic rank. The colour of the edges is proportional to the -log10 of the q-value (higher meaning more significant). The top microbial entities associated with the domain are visualised around the graph and coloured based on the taxonomic rank.

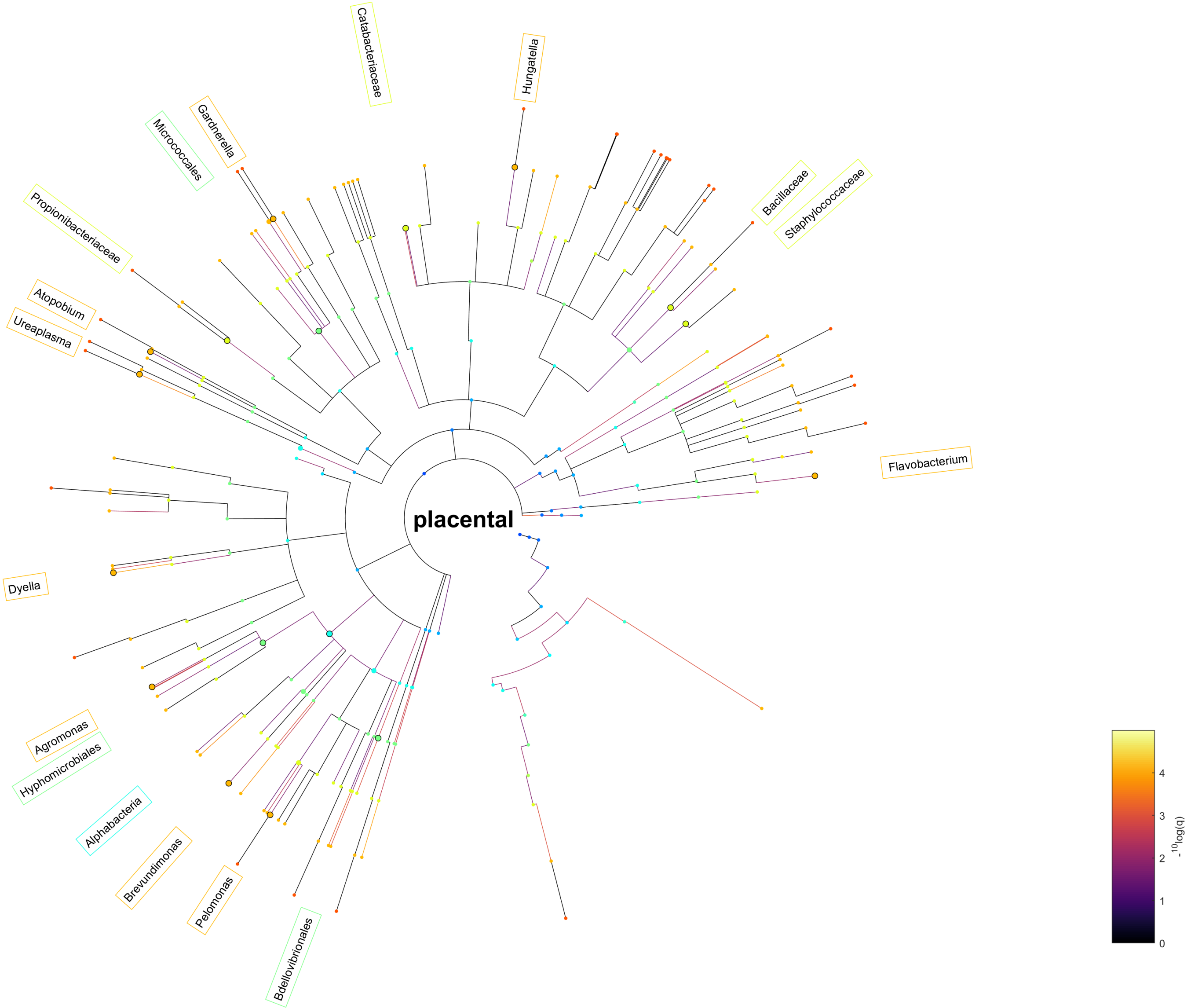


**Fig. 12.** Taxonomic tree visualisation of the placental microbiota. Colour of the nodes relate to the taxonomic rank. The colour of the edges is proportional to the -log10 of the q-value (higher meaning more significant). The top microbial entities associated with the domain are visualised around the graph and coloured based on the taxonomic rank.

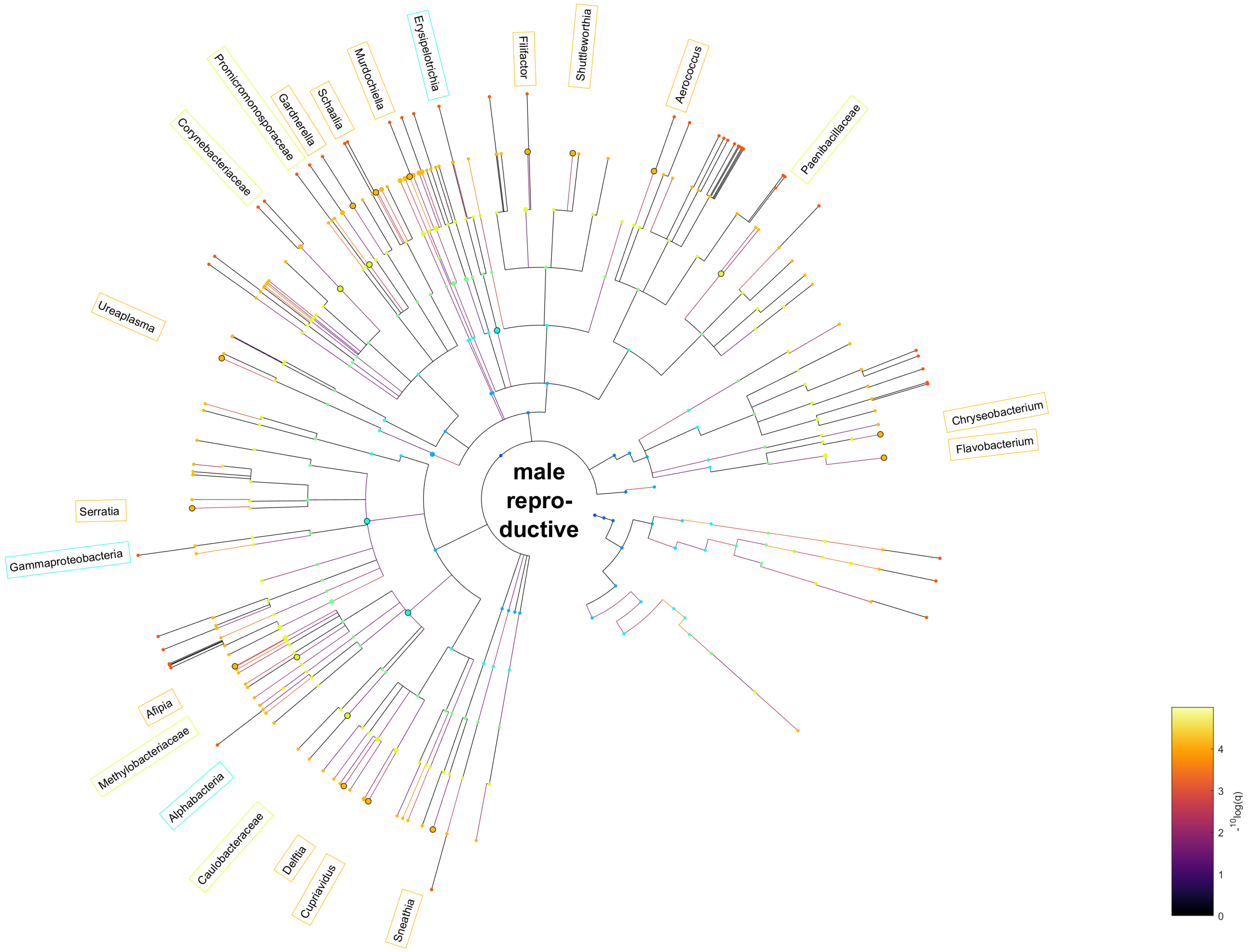


**Fig. 13.** Taxonomic tree visualisation of the male reproductive (testicle, semen, but not urinary) microbiota. Colour of the nodes relate to the taxonomic rank. The colour of the edges is proportional to the -log10 of the q-value (higher meaning more significant). The top microbial entities associated with the domain are visualised around the graph and coloured based on the taxonomic rank.

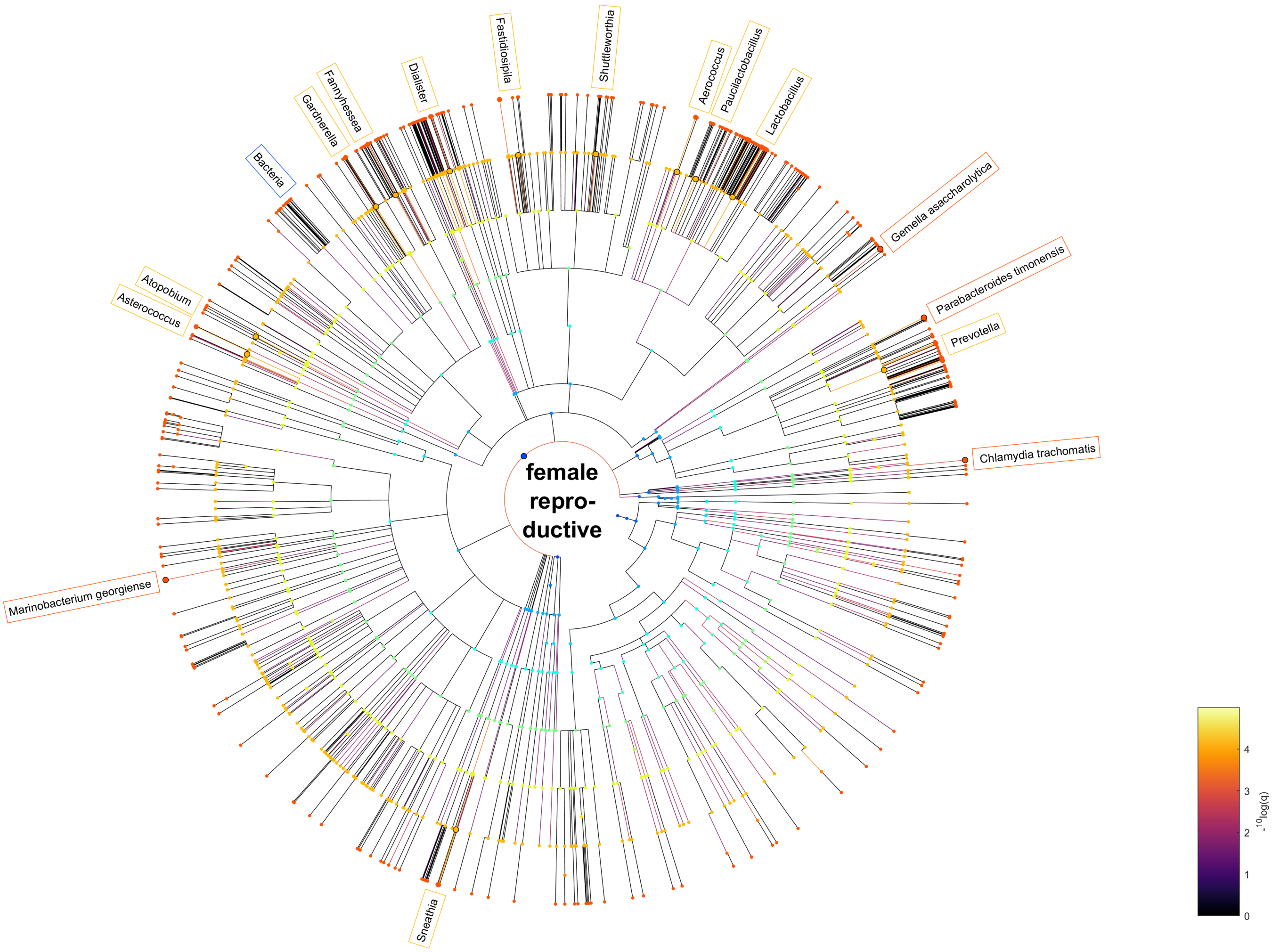


**Fig. 14.** Taxonomic tree visualisation of the female reproductive (vagina, cervix, endometrium, but not urinary) microbiota. Colour of the nodes relate to the taxonomic rank. The colour of the edges is proportional to the -log10 of the q-value (higher meaning more significant). The top microbial entities associated with the domain are visualised around the graph and coloured based on the taxonomic rank.
